# Harnessing the pangenome for genomic surveillance: *Salmonella enterica* serotype Typhi as a paradigm

**DOI:** 10.1101/2023.08.23.554479

**Authors:** Arancha Peñil-Celis, Kaitlin A Tagg, Hattie E Webb, Santiago Redondo-Salvo, Louise Francois Watkins, Luis Vielva, Chelsey Griffin, Justin Y Kim, Jason P Folster, M Pilar Garcillan-Barcia, Fernando de la Cruz

## Abstract

Public health genomic surveillance systems typically measure genome relatedness and infer molecular epidemiological relationships using chromosomal loci alone – an approximation of vertical evolution, or homology-by-descent. The accessory genome, composed of plasmids and other mobile genetic elements, reflects horizontal gene transfer and serves as an important mechanism of bacterial evolution, enabling rapid adaptation. Measuring homology in the accessory genome – homology-by-admixture – could offer important molecular epidemiological information for public health application. We applied Jaccard Index and a novel genome length distance metric to compute pangenome relatedness for the globally-important pathogen *Salmonella enterica* serotype Typhi (Typhi), and graphically express both homology-by-descent and homology-by-admixture in a reticulate network. Jaccard Index Network Analysis revealed structure in the Typhi pangenome that can be harnessed to enhance discriminatory power for surveillance, track antimicrobial resistance, and refine our understanding of homology for outbreak management and prevention. This offers a more intricate, multidimensional framework for understanding pathogen evolution.

**Significance Statement:** Bacterial relatedness is often measured and visualized using chromosomal comparison and phylogenetic trees. While valuable, this approach captures only the vertical evolutionary dimension and excludes genetic material acquired or lost through horizontal gene transfer. We present an approach for measuring and visualizing bacterial relatedness using all core and accessory genetic material and discuss the interpretation of resulting reticulate networks of bacterial genomes. In application to *Salmonella* Typhi, Jaccard Index Network Analysis revealed structure in populations of this pathogen that may be harnessed for public health applications. This approach captures both vertical and horizontal evolutionary dimensions, offering an intricate genetic framework for exploring pathogen evolution.

## Introduction

Public health surveillance systems around the world are increasingly leveraging whole-genome sequence (WGS) data and genomic methods to track and mitigate the spread of infectious diseases(*1*–*3*). Enteric foodborne surveillance has benefitted from the integration of WGS data with traditional epidemiological methods, rapidly improving outbreak detection, source attribution, and our understanding of antimicrobial resistance (AMR)(*4, 5*). Many genomic surveillance and outbreak detection systems rely on measuring core genome relatedness; for example, the United States national molecular subtyping network for foodborne disease surveillance, PulseNet USA, uses core genome multi-locus sequence typing (cgMLST) to detect single nucleotide polymorphisms (SNPs) or indels (insertions and deletions) within specific core loci(*6*). Core genome-based methods approximate vertical evolution – homology-by-descent – and offer a genetic framework for understanding epidemiological patterns(*5*). However, non-core genes– the accessory genome–reflect horizontal gene transfer (HGT), another fundamental dimension of bacterial evolution(*7*). This dimension is often overlooked in epidemiological investigations, even as phylogenetics becomes common practice.

The accessory genome is composed of plasmids, phages, and a variety of mobile genetic elements (MGEs) that exist as autonomous or chromosomally integrated molecules(*8*). It is highly variable and flexible, enabling rapid adaptation of bacterial species to new niches and environmental selection pressures(*7, 9*). Therefore, inclusion of accessory content in genomic analyses can reflect conflicting phylogenetic signals, and it is typically removed (explicitly or implicitly) to simplify genomic comparisons and epidemiological conclusions(*5, 10*). However, the accessory genome is not randomly structured, nor is it under neutral selection(*7, 9, 11, 12*); thus, relatedness in the accessory genome reflects homology-by-admixture (a term borrowed from human genetics that refers to ancestry based on gene flow from multiple sources(*13, 14*)). Measuring relatedness “by admixture” for inclusion in public-health-centered analyses could offer unparalleled epidemiological resolution(*10, 15*), particularly for enteric pathogens such as *Salmonella enterica* and *Shigella* spp. that have a pangenome dominated by accessory genome content(*16*).

Numerous tools exist for characterizing the accessory genome in WGS data, including databases for plasmids(*17*–*19*), AMR genes(*20, 21*), and MGEs(*22, 23*). This information is typically overlaid onto phylogenetic trees generated using cgMLST, whole-genome MLST (wgMLST), or SNP-based analysis to determine presence/absence patterns of known elements. While useful, relying on methods that detect only known genes, plasmids, or other genetic regions of interest limits our understanding of the contribution of the accessory genome to pathogen evolution, given it is widely uncharacterized. Additionally, overlaying accessory genome information onto a core-genome framework to understand pathogen epidemiology may mask important signals in the accessory genome, especially over short time frames where HGT is an important evolutionary mechanism(*7*). Capturing the diversity of the accessory genome and distilling it into useful metrics for public health surveillance and outbreak investigation remains challenging, both from a computational and interpretive perspective.

In this analysis, we harnessed existing tools for a novel application: to measure both homology-by-descent and homology-by-admixture. Here, we determine the contribution of the accessory genome to the evolution and epidemiology of *Salmonella enterica* serovar Typhi (herein referred to as Typhi). We employed the alignment-free genome distance estimation software BinDash(*24*) to compute exact Jaccard Index (JI) calculations as a measure of pangenome relatedness without exclusion of accessory genome content. We incorporated a novel genome length distance (GLD) metric to ensure variations generated by insertions and deletions are weighted appropriately. Typhi was chosen as a representative organism for several reasons: (a) Typhi is a globally distributed human pathogen, causing typhoid fever in an estimated 11–21 million people worldwide annually(*25*), (b) Typhi is a highly clonal organism(*26*), creating challenges for differentiation of outbreak or endemic strains using core-genome methods, (c) multidrug-resistant (MDR) and extensively drug-resistant (XDR) Typhi strains are increasingly reported, and these phenotypes are driven by MGEs such as plasmids and transposons that form part of the accessory genome(*27*–*29*). Accessory genome information will be especially valuable for understanding clonal pathogens such as Typhi and may offer improved inference of evolutionary trajectories and better resolution for public health action.

## Results

### Jaccard Index concept explanation

JI was used as a measure of similarity between all possible pairs of genomes. Each genome was transformed into its corresponding set of 21-mers and the similarity between each genome pair was calculated as the ratio of shared *k*-mers over the total number of *k*-mers (considering only once those identical between both sets and thus disregarding genome differences due to sequence repetitions). The JI value ranges from 0 to 1, where 1 indicates 100% *k*-mer similarity and 0 indicates no *k*-mers shared.

JI captures both single nucleotide polymorphisms (SNPs), either due to point mutations or recombination, and gene content differences that arise as the result of gain and loss of genetic material (indels) (Supplementary Text, Supplementary Figure 1). For two genomes differing only in *L* nucleotides, *e*.*g*., SNPs, the number of different *k*-mers will be *Lk* (the probability for two SNPs to be located in the same *k*-mer is insignificant when SNPs are not frequent). The insertion of a DNA sequence in one of the genomes will contribute *L*+*k*-1 new *k*-mers, while a deletion will render a difference of *k*-1 new *k*-mers in the deleted genome and *L*+*k*-1 in the non-deleted genome. On the other hand, the acquisition of a plasmid of size *L* will produce *L* new *k*-mers. For example, two equal genomes of 5Mb and only one of them having an 84 kb plasmid exhibit JI=0.984.

The pairwise genome similarities can be represented in an undirected network in which the nodes (genomes) are connected if the pairwise JI equals or exceeds the set JI threshold. At the initial network stage, genomes sharing any JI value greater than 0 will be linked by an edge, resulting in most genomes connected in a single component. By increasing the stringency of the JI threshold, separate connected components emerge.

### Genome length distance concept explanation

Since both SNPs and indels are reflected in JI, their individual contribution cannot be estimated from the JI value alone. Besides, in JI estimates the contribution of SNPs is disproportionate compared to that of indels. An indel of 5kb (5,000 new *k*-mers) is equivalent to 238 SNPs (238x21=4,998 new *k*-mers). As SNPs do not contribute to genome length, but indels do, a new unit of measurement, GLD (genome length distance), was defined. It uses the difference between the unique *k*-mer counts of two given genomes as a proxy for genome size variation. The lowest GLD value is 0, meaning the genomes compared do not differ in size. The theoretical upper limit to GLD depends on the size difference between the smallest and largest genomes analyzed. Two genomes with GLD=0.05 would differ in 50 kb (0.05 x 1,000,000 bp), whereas GLD=2.0 would mean a difference of at least 2 Mb between them (sequence duplications are not considered). Thus, the size of the compared genomes does not determine the GLD value, rather, it merely reflects the difference in size between them.

Pairwise GLD values can be used on top of a given JI threshold to emphasize differences in genome size (as a proxy for indels). When the GLD filter is not applied, the difference in genome size is not considered for clustering, which is equivalent to setting the GLD threshold at its upper limit. On the other extreme, when GLD=0, only genomes fulfilling the JI threshold and having equal size will be linked. Filtering the network with JI=0.983 and then applying GLD=0.05 will result in the disconnection of genomes that, despite sharing a number of *k*-mers to satisfy the JI threshold, differ in size more than 50 kb. In our dataset, the maximum GLD value is 0.764, meaning that the largest size difference between two Typhi genomes is 764 kb. GLD provides an extra layer that can be conditionally applied, depending on the dataset and resolution needed by the user.

### Population structure of Salmonella Typhi using JI network analysis

The distribution of JI values obtained from pairwise comparisons of the 2,392 genomes showed no clear valley supporting an ideal cut-off (Supplementary Fig. S2). A range of JI and GLD thresholds were assessed to generate the final network, based on the proportion of genomes assigned to JI-groups, and balancing genome differentiation with network simplification. At JI>=0.98, the relatedness of genomes within each cluster is >99.8% average nucleotide identity (ANI) (Supplementary Fig. S3). At thresholds JI=0.983 and GLD=0.05, between-group differences can be explained by indels of >50 kb in size, or by >=2,050 SNPs across the entire genome, or a mix of both (Supplementary Text). Following this approach, the optimum threshold for analyzing Typhi genomes was set at JI=0.983 and GLD=0.05, which offered a reasonable separation that enabled further molecular investigation (*i*.*e*., number and separation of groups was not overwhelming). At these thresholds, the 2,392 Typhi genomes (2,272 study genomes, and 120 RefSeq reference genomes (Supplementary Table S1)) self-organized into 17 distinct clusters according to the Louvain method, named JI-groups A-Q, with only 39/2,392 (1.6%) not assigned (singletons or JI-clusters with <5 members) (Figure 1A). JI-group A was the largest (n=1,319/2,392), with all other JI-groups represented by at least five genomes (Table 1). Three of the largest JI-groups (A, B, and C) were further divided into JI-subgroups using an increased JI threshold (Supplementary Fig. S2). JI-A subgroups A1-17 were defined at JI=0.995; JI-B subgroups B1-3 at JI=0.986; and JI-C subgroups C1-3 at JI=0.997 (Supplementary Fig. S4).

**Fig 1.**
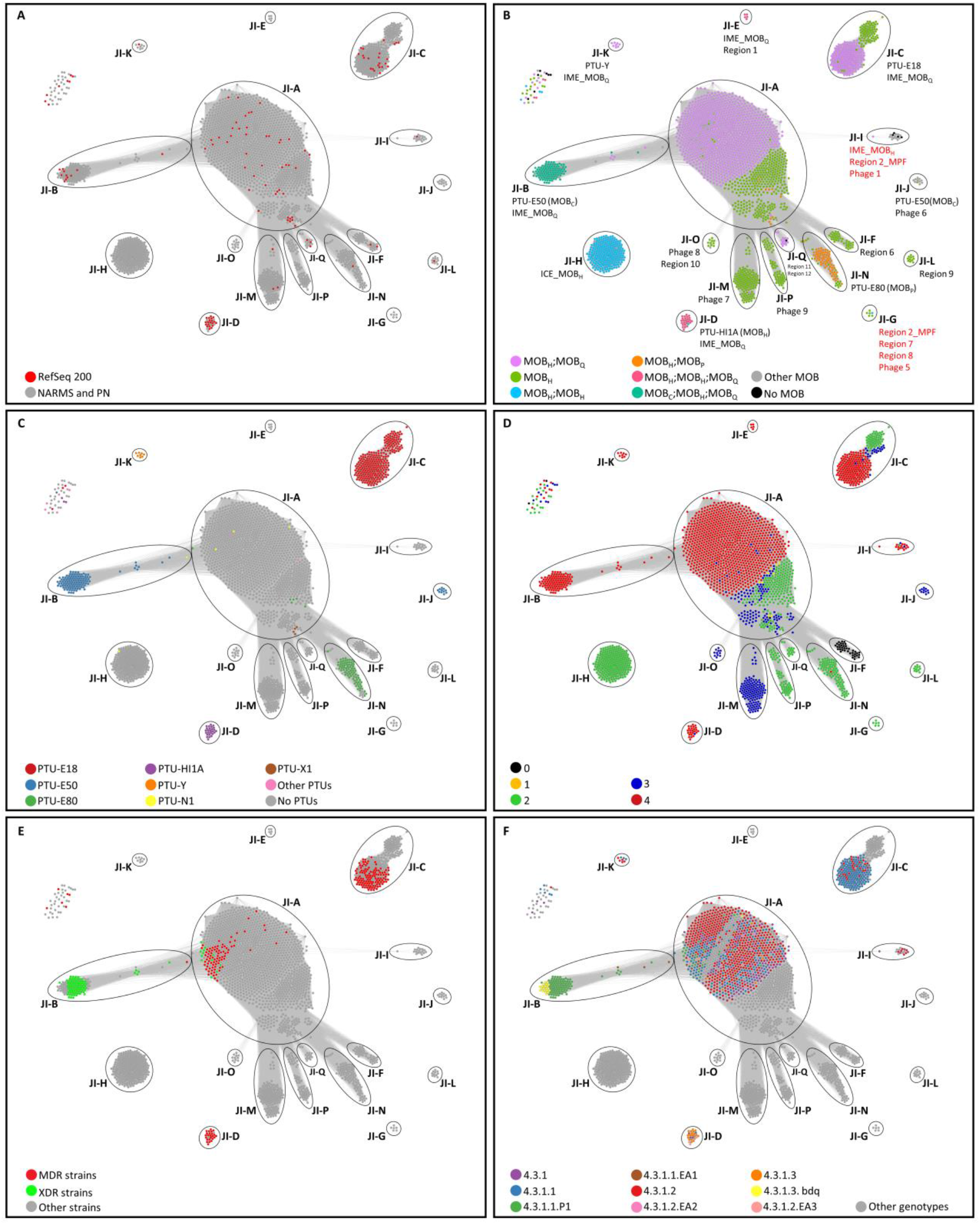
Distribution of Typhi genomes by JI. The networks contain 2,392 nodes, connected when JI>=0.983 and GLD<=0.05. Seventeen clusters (named JI-A to JI-Q) are indicated by circles. **A) Distribution of references and CDC study genomes in the JI groups**. Nodes depicted in red represent RefSeq200 genomes (references) and those in grey represent study genomes. **B) Distribution of MOB relaxases in the JI groups**. Nodes are colored according to the MOB relaxase class present in each genome. Information on the PTUs, as well as other accessory elements present in >90% of the members of a given JI group, are included in black letters when present in a given cluster or in red letters when absent (see also Supp. Fig. 4). **C) Distribution of PTUs in the JI groups**. Nodes are colored according to the PTUs present in each genome. **D) Distribution of GenoTyphi primary clades in the JI groups**. Nodes are colored according to the GenoTyphi primary clades. **E) Distribution of multidrug resistant (MDR) and extensive drug resistant (XDR) genomes**. Nodes are colored according to AMR categories. **F) Distribution of the 4.3.1 GenoTyphi genotype in the JI groups**. Nodes are colored according to the lineages and sublineages of the 4.3.1 genotype. A Gephi file containing the JI network is available at https://github.com/PenilCelis/Salmonella_Typhi_JINA.

**Table 1.** Summary of JI-group information for 2,272 US CDC and 120 RefSeq200 genomes.

| **JI-group** | **Count <sup>a</sup>** | **% <sup>b</sup>** | **GenoTyphi primary cluster** |
| --- | --- | --- | --- |
| A | 1319 | 55.1 | 0, 1, 2, 3, 4 |
| B | 114 | 4.8 | 4 |
| C | 265 | 11.1 | 2, 3, 4 |
| D | 26 | 1.1 | 3, 4 |
| E | 5 | 0.2 | 4 |
| F | 39 | 1.6 | 0 |
| G | 6 | 0.3 | 2 |
| H | 225 | 9.4 | 2 |
| I | 17 | 0.7 | 2, 3, 4 |
| J | 11 | 0.5 | 3 |
| K | 8 | 0.3 | 4 |
| L | 11 | 0.5 | 2 |
| M | 133 | 5.6 | 3 |
| N | 90 | 3.8 | 2, 4 |
| O | 11 | 0.5 | 3 |
| P | 58 | 2.4 | 2 |
| Q | 15 | 0.6 | 2 |
| Singletons | 39 | 1.6 | 0, 2, 3, 4 |

<sup>a</sup> Number of genomes present in each JI-group
<sup>b</sup> Percentage of genomes from the total dataset that belong to each JI-group

Exploring the Typhi dataset, we found that indels of accessory genome elements are responsible for most differences between JI-groups, and that JI-groups correlate with the presence/absence of specific accessory genome elements. For example, almost all (n=2,380/2,392; 99.5%) genomes carried at least one MOB relaxase gene, either on a plasmid or integrated into the chromosome (Figure 1B). Among the autonomous plasmids, recently classified in Plasmid Taxonomic Units (PTUs) (*30*), those larger than 80kb were found to contribute to the definition of some JI-groups. JI-groups B and J contain plasmids belonging to PTU-E50 (average size 90 kb); JI-group C contains PTU-E18 (average size 107 kb), JI-group D contains PTU-HI1A (average size 217 kb); and JI-group K contains PTU-Y plasmids (average size 100 kb) (Table 2, Figure 1C). Plasmids <40kb were identified in some genomes of JI-groups A, H and N; however, their relatively small size was insufficient to segregate those genomes from their “mother” JI-groups at JI=0.983 and LD=0.05. For the remaining JI-groups, the differences are mainly defined by accessory genome elements that are integrated into the chromosome (Figure 1B and Supplementary Fig. S5).

**Table 2.**
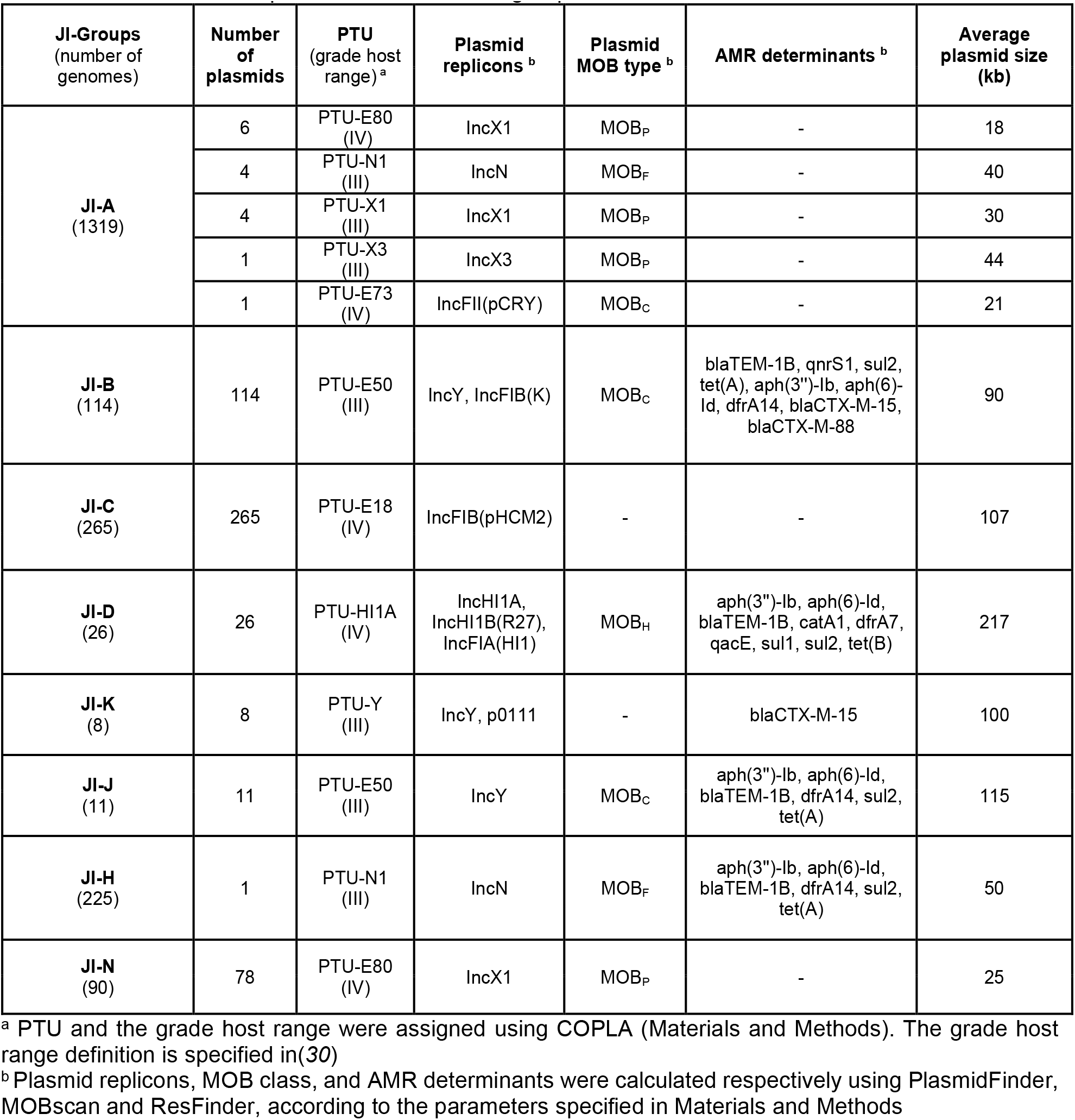
Characteristics of plasmids identified in JI-groups.

| JI-Groups<br>(number of<br>genomes) | Number<br>of<br>plasmids | PTU<br>(grade host<br>range)<sup>a</sup> | Plasmid<br>replicons<sup>b</sup> | Plasmid<br>MOB type<sup>b</sup> | AMR determinants<sup>b</sup> | Average<br>plasmid size<br>(kb) |
| --- | --- | --- | --- | --- | --- | --- |
| **JI-A**<br>(1319) | 6 | PTU-E80<br>(IV) | IncX1 | MOB<sub>P</sub> | - | 18 |
|  | 4 | PTU-N1<br>(III) | IncN | MOB<sub>F</sub> | - | 40 |
|  | 4 | PTU-X1<br>(III) | IncX1 | MOB<sub>P</sub> | - | 30 |
|  | 1 | PTU-X3<br>(III) | IncX3 | MOB<sub>P</sub> | - | 44 |
|  | 1 | PTU-E73<br>(IV) | IncFII(pCRY) | MOB<sub>C</sub> | - | 21 |
| **JI-B**<br>(114) | 114 | PTU-E50<br>(III) | IncY, IncFIB(K) | MOB<sub>C</sub> | blaTEM-1B, qnrS1, sul2,<br>tet(A), aph(3'')-Ib, aph(6)-<br>Id, dfrA14, blaCTX-M-15,<br>blaCTX-M-88 | 90 |
| **JI-C**<br>(265) | 265 | PTU-E18<br>(IV) | IncFIB(pHCM2) | - | - | 107 |
| **JI-D**<br>(26) | 26 | PTU-HI1A<br>(IV) | IncHI1A,<br>IncHI1B(R27),<br>IncFIA(HI1) | MOB<sub>H</sub> | aph(3'')-Ib, aph(6)-Id,<br>blaTEM-1B, catA1, dfrA7,<br>qacE, sul1, sul2, tet(B) | 217 |
| **JI-K**<br>(8) | 8 | PTU-Y<br>(III) | IncY, p0111 | - | blaCTX-M-15 | 100 |
| **JI-J**<br>(11) | 11 | PTU-E50<br>(III) | IncY | MOB<sub>C</sub> | aph(3'')-Ib, aph(6)-Id,<br>blaTEM-1B, dfrA14, sul2,<br>tet(A) | 115 |
| **JI-H**<br>(225) | 1 | PTU-N1<br>(III) | IncN | MOB<sub>F</sub> | aph(3'')-Ib, aph(6)-Id,<br>blaTEM-1B, dfrA14, sul2,<br>tet(A) | 50 |
| **JI-N**<br>(90) | 78 | PTU-E80<br>(IV) | IncX1 | MOB<sub>P</sub> | - | 25 |

<sup>a</sup> PTU and the grade host range were assigned using COPLA (Materials and Methods). The grade host range definition is specified in (30)
<sup>b</sup> Plasmid replicons, MOB class, and AMR determinants were calculated respectively using PlasmidFinder, MOBscan and ResFinder, according to the parameters specified in Materials and Methods

To further explore the contribution of MGEs in the JI-group clustering, an *in silico* experiment was carried out by removing them from reference genomes. PTU-E50 plasmids present in the B subgroups, and SGI11 encoded in B1 and A3 references were also eliminated. The “cured” genomes segregated from their original JI-groups and associated with the JI-A1 genomes in the network (Figure 2A). The progressive reintroduction of SGI11 (Figure 2B) and PTU-E50 sequences (Figure 2C) led to the partition of A3, B1, B2, and B3 genomes from the A1 group, rendering new clusters. In a similar experiment, large plasmids in B, C, D, J, and K reference genomes were shown to shape these JI-groups (Supplementary Fig. S6). Thus, Typhi diversification is defined predominantly by the acquisition and loss of MGEs, rather than by chromosomal point mutations.

**Fig 2.**
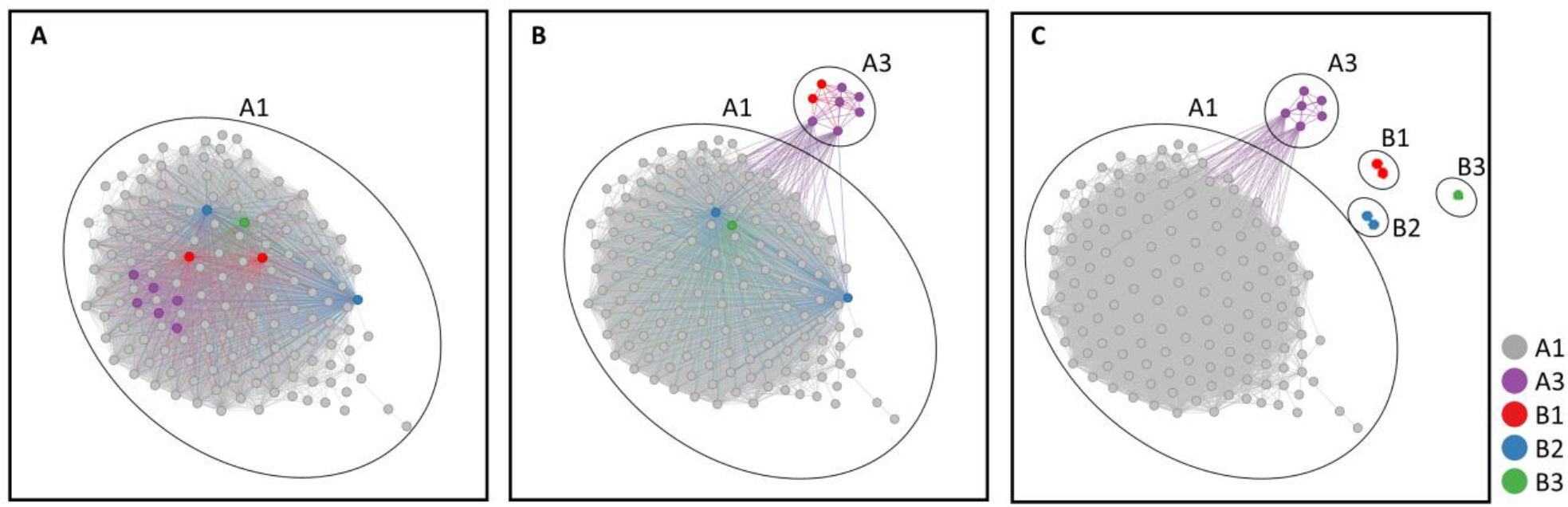
Effect of MGEs on the JI-based Typhi genome clustering. The networks contain 154 nodes, connected when JI>=0.995. Nodes are colored according to the original JI subgroup of each genome (see also Supp. Fig S3). **A) Clustering of genomes deprived of PTU-E50 and SGI11**. PTU-E50 plasmids originally present in genomes of the B1, B2, and B3 subgroups, as well as the chromosomally-inserted element SGI11, encoded also in genomes of the B1 and A3 subgroups, were removed from the genome sequences. The resulting “pruned” genomes were used to calculate pairwise genome similarities. Their reassociation with JI-A1 genomes is indicated by a circle. **B) Clustering of genomes deprived of PTU-E50**. The SGI11 elements were restituted to the A3, and B1 genomes and the network was recalculated. A3 and B1 genomes broke away from the A1 group and clustered together. **C) Clustering of genomes with SGI11 and PTU-E50**. The PTU-E50 plasmids were restituted to the B1, B2, and B3 genomes. The rebuilt network shows the emergence of distinctive clusters.

### Salmonella Typhi JI-groups are consistent across datasets

To assess whether the network obtained with US genomes is generalizable to the global population structure of Typhi, a new network was generated with a large dataset from a distinct geographic region. It included 1,606 genomes isolated in the Indian subcontinent and 136 genomes (Supplementary Table S2) from the US dataset, representative of the 17 JI-groups previously identified (Supplementary Fig. S7). The new network organized into 17 JI-groups already delimited in the US dataset (5 JI-groups are represented only by reference genomes in this network), and two new JI-groups (JI-R and JI-S, containing 10 and 11 genomes, respectively). JI-R genomes contain a PTU-X1 plasmid and three chromosomal regions enriched in phage-related genes, while JI-S members contain two plasmids (PTU-E18, PTU-HI1A) and an integrative mobilizable element. In a similar experiment, 38 Typhi genomes of the pre-antibiotic era obtained from the Murray collection(*31*) (Supplementary Table S3) were incorporated to the US genome network (Supplementary Table S1). They were distributed in groups JI-A (20 genomes), JI-F (7 genomes), JI-M (5 genomes), JI-I (1 genome), JI-Q (1 genome) and 4 isolates were singletons (Supplementary Fig. S8a), with JI-A members belonging to different subgroups (Supplementary Fig. S8b). Considering the core genome phylogeny, these ancient genomes are embedded in clades also containing recent genomes (Supplementary Fig. S8c). Finally, we generated a JI network using 1,804 globally representative Typhi genomes, which were previously used to define the GenoTyphi typing nomenclature (*32*), and 136 reference genomes from the US dataset (Supplementary Table S4 and Supplementary Fig. S9). The vast majority (1,662/1,804, 92%) of the genomes in the GenoTyphi dataset clustered in 12 of the originally defined JI groups. The rest of the genomes fell into one of eight small new JI groups (98 genomes) or were singletons (44 genomes). The application of this method to these datasets demonstrates the robustness of the JI-groups identified here, despite the caveat of data limitations.

### JI-grouping complements existing Salmonella Typhi typing methods

Typhi JI-groupings were compared to GenoTyphi genotypes(*32, 33*) for phylogenetic context. Most JI-groups (n=12/17) associated with a single subclade, clade or GenoTyphi primary clade (Table 1, Figure 1D and 1F, Supplementary Table S1), whereas JI-A, JI-C, JI-D, JI-I, and JI-N contained isolates that fell into two or more primary clades. Therefore, membership to a JI-group does not necessarily imply vertical descent (as defined by GenoTyphi). JI-A contains genomes from all four Typhi primary clades; with each primary clade mostly confined to distinct areas in the network map of JI-A (Figure 1D). JI-A, JI-B and JI-C subgroups, which were differentiated by increasing the JI threshold, were also compared to GenoTyphi genotypes (Supplementary Table S1). The majority of JI-A subgroups (n=15/17) contained genomes of a single GenoTyphi primary clade, clade or subclade (Supplementary Fig. S10, Supplementary Table S1). Similarly, all three subgroups of JI-B contain members of a single GenoTyphi subclade, while all members within each of the five subgroups of JI-C contain members of a single GenoTyphi clade or subclade (Supplementary Table S1).

### Leveraging JI network analysis to understand AMR epidemiology and evolution

*Genomic location of SGI11*. To explore patterns of AMR, acquired genes and mutations were mapped onto the JI network (JI=0.983, GLD=0.05) (Supplementary Table S1). MDR Typhi genomes were genetically defined by carriage of a complete or partial copy of the genomic island SGI11, which contains AMR genes (*bla*_TEM-1_, *catA1, aph(3′)-Ib* [*strA*], *aph(6)-Id* [*strB*], *sul1, sul2* and *dfrA7*), a mercury resistance operon, and the *qacEΔ1* gene that encodes an ethidium-bromide resistance protein(*34*). MDR genomes with SGI11 variants A and E (Supplementary Fig. S11a) belong to JI clusters JI-A, JI-C and JI-D. The genetic location of the resistance genes was consistent within each cluster – plasmid-mediated (JI-D, PTU-HIA), or chromosomal (JI-A and JI-C) (Figure 1E). XDR genomes also carry SGI11 (variants A, B, and E, Supplementary Fig. S11a) integrated in the chromosome, and were confined to JI-A and JI-B (Figure 1E), with those from JI-A collected more recently (2019–2021). SGI11 variant C and the new variant F (Supplementary Fig. S11a) were found in JI-C genomes.

SGI11 is chromosomally located in 25 reference genomes (22 in *yidA*, 3 between *cyaY* and *cyaA*), and 301 study genomes, most of them in groups JI-A, JI-B, or JI-C. In 250/301 study genomes, ISMapper identified a possible IS*1* (the flanking MGE of SGI11) insertion in *yidA*. In a core genome phylogenetic tree (Supplementary Fig. S11b), these 250 genomes were located in the same monophyletic clade, together with the reference genomes that contained SGI11 inserted in *yidA*. All SGI11 insertions in the JI-B genomes (n=88) occurred in *yidA*. Six other genomes in the same tree clade do not contain SGI11. In these cases, long-read sequencing of two of these genomes (PNUSAS224101, SAMN21040479; PNUSAS195139, SAMN18332688) confirmed that the *yidA* gene is disrupted by IS*1*, suggesting that it could be either a precursor to the SGI11 acquisition, or most likely a derivative of SGI11 excision, both probably through IS*1*-mediated recombination. It is thus possible that an insertion event of SGI11 in the *yidA* gene occurred in an ancestor of this clade and was kept in most of the descendants. For the remaining 54/301 study genomes, mostly belonging to JI-A and JI-C, ISMapper localized the IS*1* insertion site between *cyaY* and *cyaA* genes and a core genome phylogenetic tree showed that they were located in different clades that are not exclusively SGI11-containing (Supplementary Fig. S11b). This fact suggests that several unique integration events of SGI11 between *cyaY* and *cyaA* genes took place, consistent with previous analysis(*27*).

*Distribution of* bla_CTX-M-15_. The β-lactamase gene *bla*_CTX-M-15_ is a common cause of ceftriaxone resistance in XDR and non-XDR Typhi strains. A total of 109 isolates harbored *bla*_CTX-M-15_, most of them in isolates of the JI-A and JI-B groups (Figure 3), and likely mobilized by IS*Ecp1* in all cases. Of these, 93 contained *bla*_CTX-M-15_ in a plasmid whose copy number did not exceed that of the chromosome, either PTU-E50, PTU-I1 or PTU-Y. Sixteen genomes, all belonging to JI-A, contained *bla*_CTX-M-15_ in the chromosome (Figure 3). The distal position of the phylogenetic clades including these JI-A genomes in the tree suggests that the acquisition of *bla*_CTX-M-15_ occurred recently (Figure 3). Of these 16, three different sized regions of the PTU-E50 plasmid were detected in the chromosomes, likely captured and mobilized by IS*Ecp1* (Figure 4A). ISMapper identified four possible IS*Ecp1*-*bla*_CTX-M-15_ insertion sites (I-IV) (Figure 4B). The direct analysis of the *bla*_CTX-M-15_-containing contigs confirmed this location for insertion sites I-III. Long-read sequencing was performed on one genome (PNUSAS198714, SAMN18813804) containing a 17 kb region from PTU-E50 at the insertion site IV, verifying this location. Isolates with core genomes differing by fewer than 10 SNPs contain the *bla*_CTX-M-15_ gene either in a PTU-E50 plasmid or different chromosomal locations, or entirely lack it (Figure 3). This suggests that the PTU-E50-borne *bla*_CTX-M-15_ inserted in the chromosome in independent events. Genomes with a given, unique integration site, clustered together in the JI network when the JI threshold was increased (i.e., increased discrimination between genomes) (Figure 4C); however, these clusters were not distinct enough to be used for prediction of integration site from the JI network alone.

**Fig 3.**
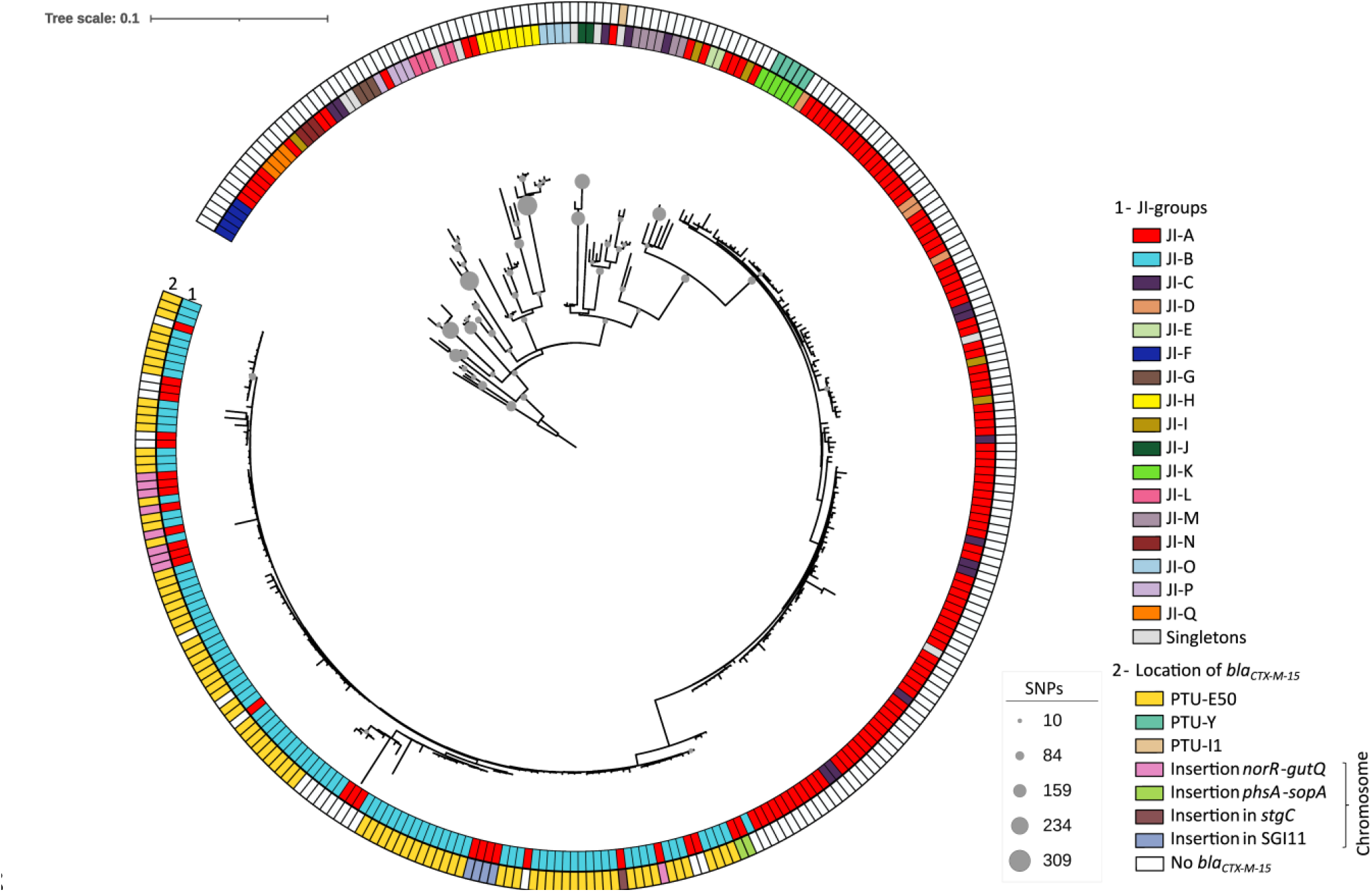
Core genome phylogeny of Typhi genomes. The cladogram includes all Typhi genomes that contain the gene *bla*_CTX-M-15_ (n=109), all genomes from JI-A3 that lack *bla*_CTX-M-15_ (n=88), all genomes from JI-B1 that lack *bla*_CTX-M-15_ (n=4), 122 representative genomes from the 17 JI groups (Supplementary Table S1), and one genome from serovar Indiana as an outgroup. Circles at the internal nodes indicate the number of SNPs distinctive of the corresponding clade. The colored rings indicate the JI-group of the corresponding genome (1), and the *bla*_CTX-M-15_ gene location (2). A phylogenetic tree of the representative genomes from the 17 JI groups is shown in Supplementary Fig. S11).

**Fig 4.**
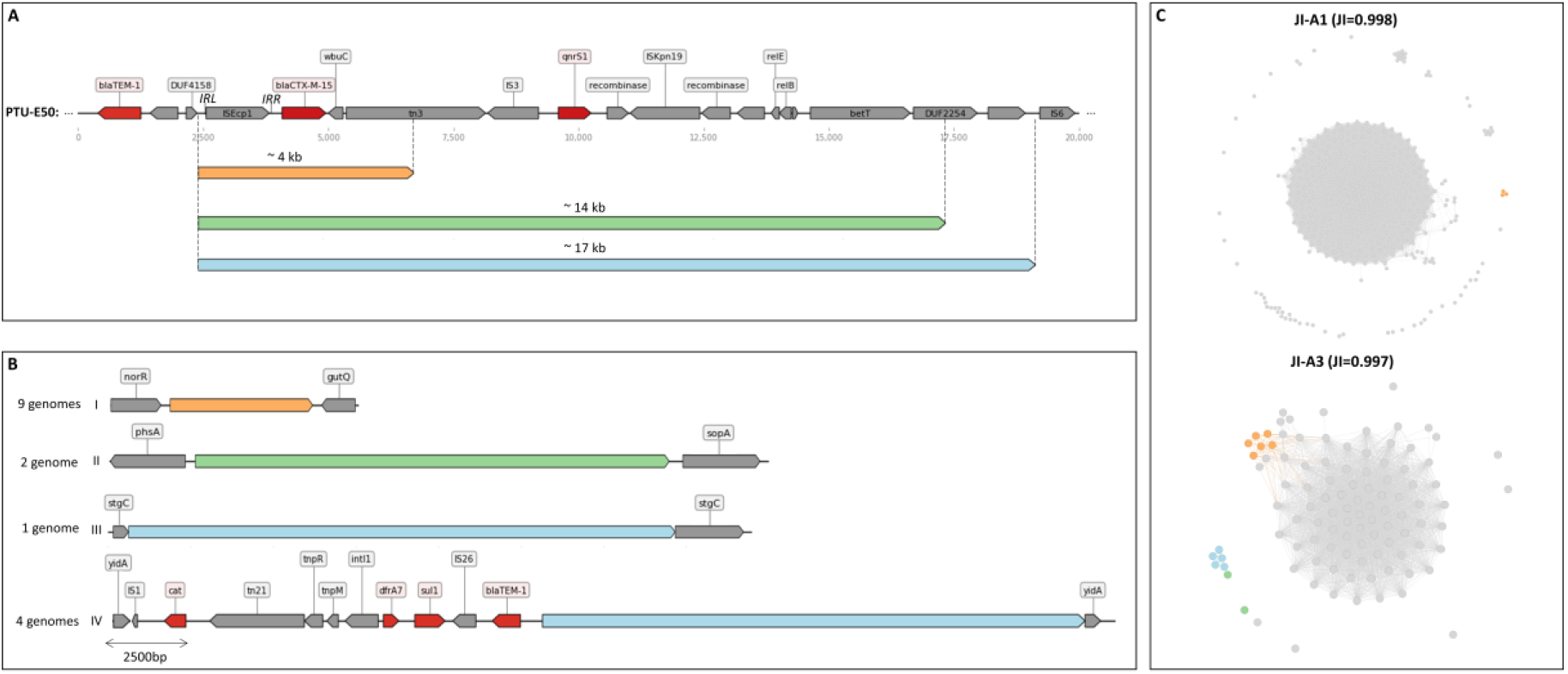
Genomic context of *bla*_CTX-M-15_. **A) The genetic vicinity of *bla***_***CTX-M-15***_ **in PTU-E50 plasmids**. The region containing the *bla*_CTX-M-15_ gene of plasmid NZ_CP046430 is depicted. Genes are represented by arrows and those encoding AMR are colored in red. Below, three arrows of different sizes, indicated by different colors represent the PTU-E50 regions that are found integrated into the chromosomes. **B) Chromosomal integration sites of the *bla***_**CTX-M-15**_**-containing regions (I-IV)**. Insertion site I locates between genes *norR* and *gutQ*; site II between genes *phsA* and *sopA*; site III interrupts gene *stgC*; site IV resides within SGI11. **C) JI networks of subgroups JI-A1** (upper panel) **and JI-A3** (lower panel). Nodes colored in orange, green, and blue indicate genomes containing the different *bla*_CTX-M-15_-encoding regions.

*Chromosomal AMR*. Chromosomal mutations in the QRDR (resistance to quinolones), and the *acrB* gene (azithromycin resistance(*35*)) were mapped onto the JI network (Supplementary Fig. S13). Genomes with one, two or three QRDR mutations tended to cluster in specific JI-groups or subgroups, however the presence of a QRDR or *acrB* gene mutation did not define the JI-groups.

### Leveraging JI network analysis to understand plasmid dynamics

Twelve different PTUs were detected in Typhi genomes in this study, predominantly from MOB_P_ and MOB_H_ classes (Table 2, Figure 1C). All PTUs were graded as host range III or higher, including those that lacked a relaxase (PTU-E18, PTU-Y); indicating their ability to colonize bacteria from different genera of the same taxonomic family. Within-serotype movement of PTUs can be seen in JI-C and JI-D, two groups defined by the presence of large (>90 kb) plasmids (PTU-E18 and PTU-HI1A, respectively) that contain genomes from different GenoTyphi primary clades (Tables 1 and 2, and Figure 1C and 1D). This suggests either a common source from which different Typhi lineages acquired similar plasmids, or between-lineage exchange of plasmids. Further, plasmid presence varied within Typhi lineages. Subclade 4.3.1.1 genomes were identified in five different JI-groups (JI-A, JI-B, JI-C, JI-D, JI-K), indicating carriage of different accessory genome content (Figure 1F) and suggesting adaptation to differing environmental selection pressures.

### Leveraging JI network analysis to understand epidemiological patterns

Epidemiological metadata was mapped onto the JI network to visualize temporal and geographical patterns (Supplementary Fig. S14). Certain JI-groups were identified throughout the entire study period from 2008–2021 (e.g., JI-A), whereas others emerged or became more prevalent in later years, such as JI-B and JI-H, in some cases due to the acquisition of a plasmid. For example, JI-B1 genomes first appeared in the US in 2018 (Supplementary Fig. S14a), and they differ from genomes in JI-A1 by the presence of PTU-E50 and SGI11 (Figure 2 and Supplementary Fig. S5). Rapid temporal “blooms” may indicate an outbreak, especially when strains share a geographic signal. In this case, JI-B1 represents the subclade responsible for an XDR Typhi outbreak that began in Pakistan in 2016(*29*). All JI-B1 members are genotype 4.3.1.1.P1, carry a PTU-E50 plasmid with *bla*_CTX-M-15_, and many are XDR. Overlay of travel data also reflects a geographic signal linked to United Nations region Asia, where Pakistan is located (Supplementary Fig. S14b). More recent genomes of genotype 4.3.1.1.P1 (2019-2021) belong to JI-A rather than JI-B1, indicating differences in accessory genome content, in this case the loss of PTU-E50. The continual dissemination of this emergent strain into different geographic locations, and over several years, likely facilitated divergence in its accessory genome. JI network visualization is able to capture this divergence, link specific genetic changes to relevant epidemiological metadata, while maintaining the ability to follow the genotype lineage initially defined as the “outbreak” strain. Following these short-term evolutionary trajectories may enable early detection of emergent strains or variants of concerning lineages linked to outbreaks.

## Discussion

Bacterial homology is used to infer epidemiological linkages in public health, using core genome targets and tree-based phylogenies to detect clusters of highly related strains that share a common epidemiological exposure(*3*). Phylogenetic trees reflect vertical evolution and assume the passage of time results mainly in the accumulation of point mutations. Typhi is a highly clonal organism, with a low mutation rate(*27*), making outbreak clusters difficult to distinguish with core-genome based methods alone. Further, emergence of MDR and XDR strains, driven by acquisition of MGEs, necessitates a focus on the accessory genome to better understand AMR dissemination. Here, we describe an approach to simultaneously analyze homology-by-descent and homology-by-admixture, enabling the incorporation of both vertical and horizontal evolutionary trajectories into our understanding of Typhi epidemiology.

JI is a common proximity measurement used to compute the similarity between two objects. It has wide use in numerous domains, such as ecology(*36, 37*), text mining(*38, 39*), and genome comparison(*24, 40*–*43*). When used to compute genome similarity, JI provides an alignment-free, quantitative assessment of similarity that can be expressed graphically in a genomic network, enabling visual integration of genomic data and available metadata. It is particularly useful for comparing and discriminating between very similar genomes (e.g., within a clonal serotype such as Typhi or other *Salmonella* serovars), because it is optimized for values well over 99.9% ANI. It takes into account both SNP and indel events (although it does not account for differences due to duplicated sequences).

The application of JI network analysis revealed that most differences between Typhi genomes can be explained by insertion and deletion of accessory genomic elements, as illustrated in Figures 1 and 2, and Supp. Fig S4. Therefore, HGT seems to be the most significant mechanism of short-term diversification in this serotype. MGEs were universally present within Typhi genomes, and each JI-group displayed a distinct accessory genome profile that corresponded to the presence or absence of particular plasmids or integrated MGEs. Thus, JI-networks reveal structure in the pangenome of Typhi, which can be harnessed for public health purposes, by enhancing discriminatory power, enabling tracking and “forecasting” of existing and novel AMR strains, and redefining our understanding of homology for outbreak management and prevention.

JI offers maximal discriminatory power by incorporation of both core and accessory genetic information. JI-groups serve as a means to readily identify the genomic structures of isolates involved in an outbreak, regardless of whether they form a new JI group or not. Thus, known outbreak lineages or endemic strains defined by core-genome methods are easily identified (Figure 1F), and within-lineage differences become obvious. For example, the XDR strain 4.3.1.1.P1, which carries a *bla*_CTX-M-15_-PTU-E50 plasmid and is responsible for a large outbreak in Pakistan(*29, 32*) clusters exclusively in JI-B1 (Supplementary Table S1). Sub-variants of this same GenoTyphi lineage can be easily identified: those that cluster with JI-A genomes (Figure 1F) have *bla*_CTX-M-15_ integrated into the chromosome and lost the PTU-E50 plasmid (Figures 3 and 4). This difference is biologically and clinically relevant because chromosomal integration of this AMR gene will likely “stabilize” the XDR phenotype in this subpopulation of Typhi genomes, with descendants of these strains all inheriting XDR. In the same way that we look to individual SNPs as unique molecular signatures for identifying subpopulations(*33*), we can exploit these accessory genome differences to enhance our discriminatory power when existing methods reach their limits. While not all molecular changes in the accessory genome will have clear biological or epidemiological significance, generating more intricate, agnostic genetic frameworks with JI analysis enables improved stratification of Typhi populations.

AMR emergence in Typhi (and enteric bacteria more generally) is governed by the movement of plasmids and other MGEs. Thus, any attempt to understand the evolution of MDR and XDR strains, or the emergence of novel AMR strains, requires knowledge of accessory genome dynamics. In Typhi, JI network analysis revealed intra- and inter-species plasmid dynamics that likely influence the evolution and emergence of AMR. First, two plasmid families associated with AMR (PTU-E50 and PTU-Y, both carrying *bla*_CTX-M-15_) displayed enough within-PTU variance to suggest plasmids of these PTUs were acquired by Typhi more than once (Supplementary Fig. S15). The data suggest that acquisition of AMR plasmids is likely a continuous occurrence in Typhi. Secondly, the abundance of chromosomally integrated MGE (Figure 1B) highlights the potential for integration of AMR regions on MGE; thus, we should expect to see continual “stabilization” of AMR phenotypes in the chromosome, which may create opportunities for new AMR plasmids to enter and establish themselves. Third, the main PTUs identified in Typhi have host range III or above (Table 2)(*30*), indicating that they exchange genetic information with different bacterial genera or families. This means the emergence and epidemiology of AMR in Typhi is not restricted by the “resistome” or “plasmidome” of *Salmonella*, but is subject to gene exchange networks that extend well beyond the species barrier(*30*).

The increasing occurrence of “plasmid outbreaks”, characterized not by a clonal strain or single species but by the presence of a shared plasmid(*44, 45*), challenges the notion of homology-by-descent in public health spaces. Further, the potential for a single-species, multi-lineage outbreaks defined by a shared MGE is also possible, at least for Typhi, given the co-occurrence of diverse strains in a relatively small geographic area(*46*). Acquisition of a common MGE by such diverse strains, that is, homology-by-admixture, may reflect adaptive evolution to a common niche or exposure, and thus exhibit epidemiological relevance. JI network analysis allows the capture and visualization of this evolutionary dimension within a particular serotype.

A potential alternative to infer genealogy from large-scale genome datasets based on alignment-free methods, which exploits core and accessory genomic variation for genome comparison, is PopPUNK (*43*). There is currently no publically available database or refined model for *S. enterica*. Even so, PopPUNK is especially suited to differentiate clonal complexes or sequence types from a given species, rather than to assess the differences within them. Our objective in this study was to utilize JI to highlight variations within a highly clonal *S. enterica* serotype. When applied to our dataset, the number of clusters and networks obtained using PopPUNK varied based on different parameters. Even when using identical parameters during pipeline execution, the generated networks were not robust, probably due to the high clonality of our dataset.

Given homology can arise in a reticulate fashion, capturing phylogenetic patterns (SNPs) and MGE dynamics in a network, such as that generated with JI, encompasses both horizontal and vertical evolutionary dynamics, which are important for understanding epidemiological patterns. Incorporating this notion into our framework of interpretation offers a more intricate, multidimensional understanding of Typhi evolution, which can be extended to other pathogens of concern. Attempts to better characterize the “what”, can elucidate the “how”, which is foundational to understanding the “why” of pathogen evolution(*47*).

In summary, we describe JI network analysis as a pangenome analysis approach to characterize genomes at a more granular level. Knowing that current core-genome based approaches exclude a proportion of genetic material, we discuss ways in which JI network analysis could benefit surveillance systems by providing higher molecular resolution to differentiate closely-related genomes, and a way to identify genomes that share identical accessory elements, which may indicate a common reservoir. We do not propose this method as a replacement to existing core-genome based public health surveillance systems, but as a complementary analysis approach to further stratify clonal serotypes. In a public health setting, this has the potential to focus on the most epidemiologically relevant genetic events, and early detection of strains of concern.

## Materials and Methods

### Isolate collection and metadata

Typhi is a nationally notifiable disease in the United States (https://ndc.services.cdc.gov/case-definitions/salmonella-typhi-infection-2019/). The Centers for Disease Control and Prevention (CDC) request state and participating local public health laboratories (PHL) (https://www.cdc.gov/narms/index.html) to submit all Typhi isolates that they receive from clinical laboratories to the National Antimicrobial Resistance Monitoring System (NARMS). Since 2016, NARMS and PulseNet USA, an enteric disease surveillance network of state and local PHL, have routinely performed WGS on Typhi isolates(*6*). CDC’s National Typhoid and Paratyphoid Fever Surveillance system collects metadata on all Typhi cases reported to PHL, including history of international travel in the 30 days before illness onset (https://www.cdc.gov/typhoid-fever/surveillance.html).

### Whole genome sequencing

WGS data was available for 2,272 Typhi isolates collected from January 1, 2008, through September 30, 2021 (Supplementary Table S1). For years prior to routine WGS (2008–2015), all Typhi isolates in the PulseNet national database with WGS data available were included (n=68); these isolates represent a small proportion of total isolates from this time period. For years 2016–2018, all Typhi isolates sent to NARMS for WGS were included (n=1,343), which is representative of US Typhi cases reported to CDC for these years. Due to logistics and delays in shipping for NARMS surveillance isolates in recent years, the 2019–2021 time period is represented by Typhi isolates in PulseNet with WGS data available (n=861), with expected underreporting due to SARS-CoV-2 pandemic-related factors. WGS performed through NARMS and PulseNet followed standard operating procedures for the Illumina Miseq platform (https://www.cdc.gov/pulsenet/pdf/PNL38-WGS-on-MiSeq-508.pdf). Reads with a base call quality score ≥28, and coverage ≥40 x were assembled using shovill v.1.0.9 (https://github.com/tseemann/shovill); and contigs with coverage below 10% average genome coverage were excluded from final assemblies.

Long-read sequencing was performed on select isolates as previously described(*48*). Corresponding Illumina short reads were generated from the same DNA extraction; libraries were prepared using the Illumina DNA Flex preparation kit per the PulseNet protocol (https://www.cdc.gov/pulsenet/pdf/PNL35_DNA_Flex_Protocol_Lib_Prep-508.pdf) and sequenced on the Illumina MiSeq platform as described above. Hybrid assemblies were generated as previously described(*48*) and uploaded to National Center for Biotechnology Information (NCBI).

### Molecular subtyping and characterization

Typhi study genomes were typed using the updated GenoTyphi scheme(*32, 33*) (https://github.com/katholt/genotyphi). AMR determinants were detected using staramr, v.0.4.0(*49*), which employs the ResFinder database (updated 30JUL2020; 90% identity, 50% gene coverage) and the *Salmonella* spp. PointFinder scheme (updated 30AUG2019). Accessory genome elements were detected using a database adapted from PlasmidFinder(*17*) (90% identity, 60% gene coverage) for plasmid replicons, MOBscan(*18*) for conjugative relaxases, CONJScan(*50*) for conjugative systems, and COPLA(*19*) to assign plasmids to a given PTU(*30*). Reconstruction of plasmids from Illumina reads was performed using PLACNETw(*51*).

MDR was defined as the presence of genes conferring resistance to ampicillin, chloramphenicol and trimethoprim-sulfamethoxazole, which are typically acquired within an IS*1*-mediated composite transposon, either on a plasmid or integrated into the chromosome as SGI11 (*Salmonella* genomic island)(*32, 33, 52*). XDR was defined as MDR with the addition of a ciprofloxacin resistance mechanism (quinolone resistance determining region [QRDR] mutation and/or plasmid-mediated quinolone resistance [PMQR] gene), and a ceftriaxone resistance gene (typically *bla*_CTX-M-15_)(*29, 53*).

Chromosomal integration events were detected using the typing mode of ISMapper(*54*) to identify acquisition of an insertion sequence (IS) relative to a reference chromosome. To detect the integration sites of *bla*_CTX-M-15_, its mobilizer, IS*Ecp1*, was used as a bait against a reference chromosome (Typhi 311189_291186, NZ_CP029894.1). Integration of SGI11 was detected using IS*1* as a bait element, and Typhi CT18 as the reference chromosome (NC_003198).

### Additional genomes

One-hundred twenty Typhi reference genomes from NCBI RefSeq200 database (accessed on 14MAY2020) were included in the analysis, collected between 1916 and 2019 (Supplementary Table S1). A dataset for comparative analysis against the US dataset was generated using all Typhi genomes isolated in the Indian subcontinent available in Pathogenwatch(*55*) (n=1,606) (accessed on 22MAR2021). Specifically, this dataset included genomes linked to Bangladesh (n=637), India (n=487), Nepal (n=318), Pakistan (n=158), and Sri Lanka (n=3), or a combination of these countries (n=3) (Supplementary Table S2).

Thirty-eight Typhi genomes from the Murray collection(*31*) available at the European Nucleotide Archive at (https://www.ebi.ac.uk/ena/browser/view/PRJEB3255) (Supplementary Table S3) and a database of 1,804 globally representative *S*. Typhi genomes used to develop the GenoTyphi typing scheme (*32*) available at Pathogenwatch (https://pathogen.watch/genomes/all?collection=nti046ubbs7t-wong-et-al-2015&genusId=590) (Supplementary Table S4) were included in a comparative analysis against the US dataset. Molecular subtyping and characterization of reference genomes was performed as above.

### Jaccard Index and Length Distance analysis

The exact Jaccard Index (JI) was used as a measure of similarity between all genome pairs. First, the complete assembly of each genome was converted into a set of *k*-mers. JI was calculated as the ratio of shared *k*-mers over the total number of different *k*-mers between the two sets (shared *k*-mers, SNP *k*-mers (i.e., *k*-mers differing by just one bp), and indel *k*-mers (i.e., *k*-mers different in both datasets)). BinDash(*24*) was used to calculate JI, using parameters minhashtype=-1 (to compute the exact JI) and *k*-mer length (*k*)*=*21 (this latter as previously defined as optimum in (*40*)). The formula to calculate JI between genomes A and B (1) is defined as the size of the intersection divided by the size of the union of the two *k*-mer sets of genomes A and B.

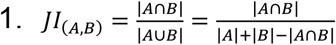

To calculate the contribution of SNPs and indels, the JI formula can be rewritten into two specific formulas, respectively (2) and (3), using parameters *k* (*k*-mer length), *L* (number of SNPs or the inserted region in size bp), and *N* (total number of genome *k*-mers of the reference genome) (Supplementary Text). Both formulas provide a rule of thumb estimation on the specific contributions of SNPs and indels to the JI value.

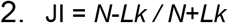

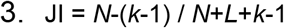

Genome length was estimated from the number of unique *k*-mers in a genome (*S*). The upper *k*-mer length limit in Jellyfish v.2.2.6(*56*) (*k*=27) was used to generate *k*-mers from each genome sequence because with greater *k*-mer length, the probability of having repeated *k*-mers by chance in a genome is lower and the genome length estimation is more accurate. *S* was computed by counting the occurrences of identical *k*-mers only once, that is, unique *k*-mers. To obtain a relative measure of genome size, the unique *k*-mer count is divided by 1 million base pairs (*S* / 1000000). For every genome pair (A and B), the difference between their unique *k*-mer counts is recorded as a length distance (GLD) value (4).

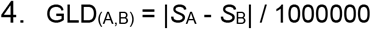

Taking into account that contig ends affect *k*-mer count, a correction described by formula (5) was applied in draft genomes, considering *S, k*, and the number of contigs of the assembly (*C*).

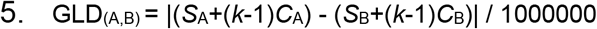

The adjacency matrix of pairwise genome similarities generated by BinDash was used to build an undirected network. Gephi (https://gephi.org/(*57*)) was used to visualize the network, applying the ForceAtlas2 algorithm for the layout. The network nodes, representing genomes, were colored according to metadata and genetic determinants of interest. Edges between nodes are represented whenever the corresponding JI or GLD value is equal to or higher than the user-defined threshold. A range of JI thresholds for a given application needs to be assessed to define the final components to study, referred to as JI-groups. This depends on the specific study population and question pursued, but it is recommended to minimize complexity by setting a threshold that will result in a manageable number of JI-groups (i.e., number of clusters should not exceed the square root of the number of genomes); to group the greatest number of genomes possible; and to factor in congruence with genetic determinants of interest, if available. The Louvain method (*58*), implemented in Gephi, was used to define the JI-groups by using resolution 1.5. Once the main JI-groups are defined, they can be further dissected in several subgroups within the network using a more stringent JI (Supplementary Figure S1) and the same community detection algorithm was used.

Distinct differences in indels, including MGEs or accessory genome regions, between JI-groups were detected using BLAST (v. 2.6.0+) by comparing reference genomes from each JI-group (Supplementary Table S1). For those JI-groups that did not contain a reference genome, a genome was reconstructed using PLACNETw(*51*). Plasmid presence was also detected using PlasmidSeeker(*59*). The BLAST searches between all possible reference pairs from different JI-groups enable the detection of regions present in one genome of the pair and absent in the other, and an estimation of their expected size can be obtained by clearing *L* from formula (3). If the size of the genome-specific regions targeted by BLAST is similar to *L*, it can be assumed that genome differences are due mainly to indels. Otherwise, SNPs or a mix of SNPs and indels account for the differences.

### Phylogeny reconstruction

kSNP 3.0(*60*) was used to identify SNPs in WGS data (complete, assembled, and raw short-read data) using *k*-mers=19. This optimal *k* was chosen with the kSNP tool Kchooser. SNP-based trees were reconstructed by maximum parsimony using the core-SNPs detected (option -core). All the trees generated in this study were visualized with iTol v6(*61*).

### Plasmid copy number calculation

The presence of specific plasmids and their average plasmid copy number (PCN) was estimated from the sequence read files of 1,836 Typhi isolates using PlasmidSeeker(*57*), including 1,157 genomes from JI-A, 97 from JI-B, 152 from JI-C, 5 from JI-D, 5 from JI-E, 7 from JI-F, 6 from JI-G, 158 from JI-H, 9 from JI-I, 10 from JI-J, 7 from JI-K, 4 from JI-L, 117 from JI-M, 75 from JI-N, and 28 singletons. PlasmidSeeker was executed with default parameters and a plasmid database of 1,064 plasmids from RefSeq200 (1,011 from *Salmonella* spp., and 153 belonging to PTU-E50, PTU-Y, PTU-E7, and PTU-E80 from Enterobacterales).

## Acknowledgments

We acknowledge the state and local public health laboratories that participate in PulseNet and the National Antimicrobial Resistance Monitoring System (NARMS). We thank the laboratorians and epidemiologists that offered helpful review and critique.

This work was supported by:

The Centers for Disease Control and Prevention, Contracts No. 75D30119C06679 and 75D30121C11978 (FdlC)

Spanish Ministry of Science and Innovation, MCIN/AEI/10.13039/501100011033 PID2020-117923GB-I00 (FdlC and MPG-B).

Spanish Ministry of Economy, Industry and Competitiveness, DI-17-09164 (SR-S).

