## Supplementary material for "Harnessing the pangenome for genomic surveillance: *Salmonella enterica* serotype Typhi as a paradigm": Supplementary Materials_Penil-Celis_Tagg_2023.pdf

**This PDF file includes:**

Supporting Information Text
Supplementary Methods
Figures S1 to S15
Legends for Datasets S1 to S4
SI References

**Other supporting materials for this manuscript include the following:**

Datasets S1 to S4

**Supporting Information Text**

Additional explanatory information on Jaccard Index definition

The Jaccard Index (JI), or Jaccard similarity coefficient, is a statistic defined as the ratio of the size of the intersection of two datasets (A and B) over the size of their union, as shown by Eq. 1:

$$JI_{(A,B)} = \frac{|A \cap B|}{|A \cup B|} = \frac{|A \cap B|}{|A| + |B| - |A \cap B|}$$

**Eq. 1**

Therefore, $0 \leq JI_{(A,B)} \geq 1$, where values closer to 1 indicate great overlap, that is, higher similarity between A and B datasets. JI is a symmetrical measure, that is, $JI_{(A,B)} = JI_{(B,A)}$. A closely related notion, the Jaccard distance, measures the dissimilarity of two datasets, as defined by Eq. 2:

$$d_{JI(A,B)} = 1 - JI_{(A,B)} = \frac{|A \cup B| - |A \cap B|}{|A \cup B|} = \frac{|A \,\Delta\, B|}{|A \cup B|}$$

**Eq. 2**

Applying JI to compare genomic sequences can be understood, at first glance, as a simple exercise of counting k-mers in both sequences (a k-mer is any contiguous sequence of *k* nucleotides present in a DNA sequence). Two characteristics of DNA can affect JI estimation. First, the molecular topology of DNA sequences, whether they are linear or circular, affects k-mer computation in contig edges (as shown in Eq. 3):

$$\#\text{ of kmers} = \begin{cases} N, & if\ circular\ contig\ of\ size\ N \\ N - k + 1, & if\ linear\ contig\ of\ size\ N \end{cases}$$

**Eq. 3**

Second, the double stranded nature of DNA sequences, which makes it impossible to know each contig strand in a draft genome. To overcome this problem, k-mers have to be extracted from both strands of the sequences (the one corresponding to the original contig and the one corresponding to its reverse complement). The other approach is to use canonical k-mers, which are defined as those that come earlier in alphabetical order of a sequence and their reverse complement. Using canonical k-mers the strand direction of the assembled contigs can simply be ignored and Eq. 3 is valid. A drawback of using canonical k-mers is that the k-mer space is halved and, thus, the probability of duplicated k-mers occurring by random chance for a specific *k* value is doubled. To avoid this problem a large enough *k* value must be chosen.

Influence of SNPs in the Jaccard Index

Let A and B be two genomes of equal size *N*. If a mutation is introduced in a position of sequence B, the number of different canonical k-mers between the two sequences will be *k*. Therefore, the Jaccard Index between both sequences is obtained from Eq. 4:

$$JI_{(A,B)} = \frac{|A \cap B|}{|A \cup B|} = \frac{N - k}{N + k}$$

**Eq. 4**

For the multiple SNPs case, if the number of SNPs (*L*) is very small relative to the genome size (*N*), and assuming that SNPs are produced following a Poisson distribution, the probability for two SNPs to be located in the same k-mer (i.e. closer than *k* base pairs) can be disregarded, and JI could be calculated with the formula:

$$JI_{(A,B)} = \frac{|A \cap B|}{|A \cup B|} = \frac{N - Lk}{N + Lk}$$

**Eq. 5**

Eq. 5 can be illustrated with a simple example: genomes A and B are two identical circular sequences of length *N*=22 (thus, the length is defined as the total number of k-mers). Three SNPs (*L*=3) were introduced in the sequence of genome B (Figure S1a). The number of different k-mers between the two sequences will be *Lk*. Therefore, the Jaccard Index between both sequences is:

$$JI_{(A,B)} = \frac{N - Lk}{N + Lk} = 0.19$$

In the case of SNP hotspots, the influence in k-mers for each SNP is less than *k*. Figure S1b illustrates this situation.

The following table shows the influence of increasing amounts of SNPs in the Jaccard Index for typical values in real cases of bacterial genomes comparisons.

**Discrimination of SNPs by using the Jaccard Index with k=21 and k=31 in a sequence of length N=5x10$^6$.**

| SNPS (*L*) | SNPS/MB | *D* = *L* / *N* [A] | *%ID* = 1 - *D* [B] | $JI_{21}$ [C] | $JI_{31}$ [D] |
|---|---|---|---|---|---|
| 50 | 10 | 0.00001 | 0.99999 | 0.99958 | 0.99938 |
| 100 | 20 | 0.00002 | 0.99998 | 0.99916 | 0.99876 |
| 500 | 100 | 0.0001 | 0.9999 | 0.99581 | 0.99383 |
| 1000 | 200 | 0.0002 | 0.9998 | 0.99165 | 0.98771 |
| 2050 | 410 | 0.00041 | 0.99959 | 0.983 | 0.97506 |
| 5000 | 1000 | 0.001 | 0.999 | 0.95928 | 0.94076 |
| 5880 | 1176 | 0.00118 | 0.99882 | 0.95237 | 0.93088 |
| 10000 | 2000 | 0.002 | 0.998 | 0.92099 | 0.88658 |
| 50000 | 10000 | 0.01 | 0.99 | 0.6815 | 0.57909 |
| 250000 | 50000 | 0.05 | 0.95 | 0.21208 | 0.11872 |

[A] Sequence dissimilarity. [B] Percentage of sequence identity. [C] Jaccard Index computed with *k* = 21. [D] JI computed with *k* = 31. In blue, the row corresponding to 5,880 SNPs, which results in the same Jaccard Index as a 250 kb insertion in a 5 Mb genome (see Figure S1f). In green, the row corresponding to JI=0.983, the threshold used to sparsify the Typhi network.

As the number of SNPs gets larger the probability of two SNPs occurring in the same k-mer cannot be dismissed, thus Eq. 5 is not a good JI estimator. In general, for any abundance of SNPs JI can be calculated by using Eq. 4 in (*1*), which estimates JI as a function of the sequence dissimilarity (*D* = *L* / *N*):

$$JI_{(A,B)} = 1/(2e^{kD} - 1) = 1/\left(2e^{kL/N} - 1\right)$$

**Eq. 6**

Influence of Indels in the Jaccard Index

The acquisition of a plasmid of size *L* by a genome A of size *N* is simply a particular case of sequence insertion not affected by the k-mer length, and its contribution to the JI is defined in Eq. 7:

$$JI_{(A,B)} = \frac{|A \cap B|}{|A \cup B|} = \frac{N}{N + L} = 1 - \frac{L}{N + L}$$

**Eq. 7**

Sequence insertion in a genome is slightly more complicated. Let A be a genome with a sequence of length *N*, and B a genome equal to A, except for an insertion of length *L* (size of genome B = *N* + *L*). Barring duplicated k-mers because of random k-mer repetitions or duplication events, it can be assumed that B contains all common k-mers with A plus *L* + *k* – 1 new ones (see Figure S1c). At the same time, in genome A, *k* – 1 k-mers around the insertion point will be specific for this genome. Therefore, the Jaccard Index between sequences A and B will be:

$$JI_{(A,B)} = \frac{|A \cap B|}{|A \cup B|} = \frac{N - (k - 1)}{N + L + k - 1} = 1 - \frac{L + 2k - 2}{N + L + k - 1} » 1 - \frac{L}{N + L}$$

**Eq. 8**

When *k* << *L* k-mer size effect on Eq. 8 can be neglected and thus Eq. 8 is transformed into Eq. 7. Eq. 8 can be illustrated with a simple example: let genome A be a circular genome of length *N*=22, and B a genome equal to A, except for a 7bp insertion (*L*=7). The Jaccard Index between sequences A and B using k-mers of length *k*=3 will be:

$$JI_{(A,B)} = \frac{N-k+1}{N+L+k-1} = 0.55$$

For the plasmid loss case, the formula (Eq. 9) is slightly different from the plasmid gain case shown in Eq. 7:

$$JI_{(A,B)} = \frac{|A \cap B|}{|A \cup B|} = \frac{N-L}{N} = 1 - \frac{L}{N}$$

**Eq. 9**

Eq. 10 allows JI calculation when a sequence is deleted in one of the genomes (see Figure S1d). When *k* << *L*, the Eq. 10 result tends to be equal to that obtained by applying Eq. 9.

$$JI_{(A,B)} = \frac{|A \cap B|}{|A \cup B|} = \frac{N-(L+k-1)}{N+k-1} = 1 - \frac{L+2k-2}{N+k-1} \text{ » } 1 - \frac{L}{N}$$

**Eq. 10**

Eq. 9 can be illustrated with the following example: let genome A be a circular sequence of length *N*=22 and B a genome identical to A except for a 7bp deletion (*L*=7). The Jaccard Index between both sequences can be estimated as follows:

$$JI_{(A,B)} = \frac{N-L-k+1}{N+k-1} = 0.42$$

<u>Influence of sequence replacement in JI</u>

A special case is the substitution of a DNA stretch of *L* bp (see Figure S1e). This case includes the occurrence of a cluster of closely located SNPs (less than *k* nucleotides of distance between each successive pair). The formula to calculate JI will be:

$$JI_{(A,B)} = \frac{|A \cap B|}{|A \cup B|} = \frac{N-\left(L+(k-1)\right)}{N-(L+k-1)+(L+k-1)+(L+k-1)} = \frac{N-L+k-1}{N+L+k-1} \approx \frac{N-L}{N+L}$$

**Eq. 11**

Eq. 11 can be illustrated with the following example: let A and B be circular genomes of length *N*=22 that differ in a sequence stretch of 7bp (*L*=7). The Jaccard Index between sequences A and B will be:

$$JI_{(A,B)} = \frac{N-L+k-1}{N+L+k-1} = 0.58$$

<u>JI vs sequence identity</u>

A graphical representation of the influence of a variety of genome differences in the Jaccard Index is shown in Figure S1f.

**Supplementary Methods**

Average nucleotide identity calculations

FastANI *(2)* was used to estimate the within JI-group genome similarity, using the 2,392 genomes of the US CDC dataset.

Phylogeny reconstruction of PTU-E50 and PTU-Y plasmids

The *pangenome* and *corepers* modules of PanACoTA *(3)* were used to detect the core genome of plasmids belonging to either PTU-E50 or PTU-Y. The *align* module was used to build the core genome alignment, which was used in turn to reconstruct a ML phylogeny with IQ-TREE 2 *(4)*, using the HKY+F model for PTU-E50 and TIM3e+R3 model for PTU-Y. ModelFinder *(5)* was used to identify the best substitution model according to the Bayesian Information Criterion. Branch support was estimated by performing 1000 ultrafast bootstrap approximation replicates *(6).*

Plasmid ORFeome analysis

AcCNET *(7)* was used to build the PTU-E50 ORFeome network. Homologous protein clusters were generated using kClust (*8)*, with >95% protein identity, >80% alignment coverage and clustering E-value < 1E-14. All edges were assigned equal weights.

Plasmid comparative genome analysis

The comparative genomic analysis of the different plasmids found in JI-B and in JI-K was carried out with Blast Ring Image Generator (BRIG) (*9*) using default parameters. It represent the sequence similarity between each query with the choosen reference. Nucleotide identity ≥50% is indicated by a band colored according to the color code, with darker shading corresponding to higher sequence match. Blank regions indicate <50% nucleotide identity. Predicted coding sequences (CDS) of the reference genome are indicated in the outermost black ring.

### Supplementary Figures

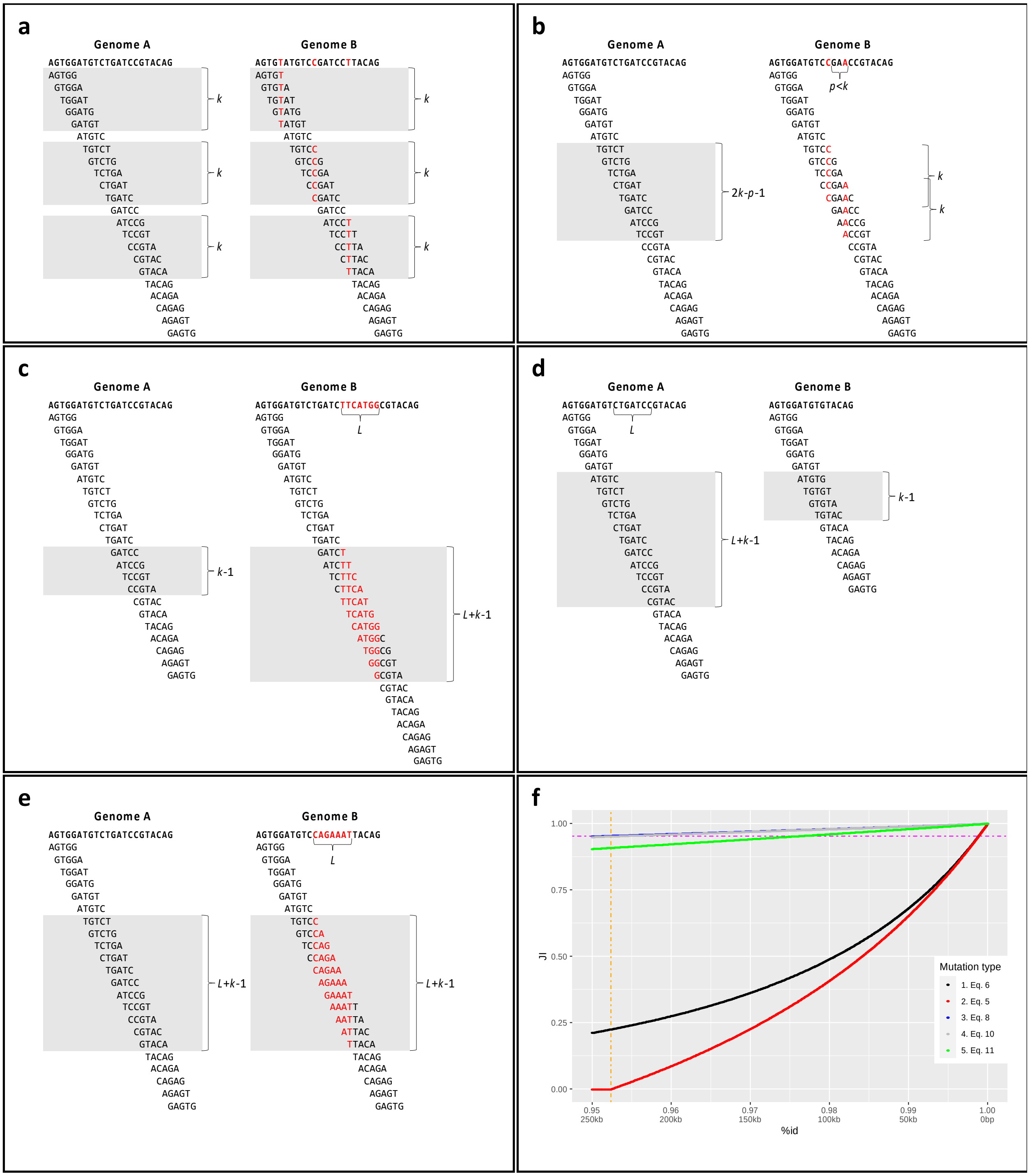


**Figure S1: Influence of SNPs and indels in JI. (a) Influence of SNPs between two sequences.** The sequence of each genome is split into k-mers of length *k*=5. Both genomes differ in *L*=3 SNPs (colored in red in genome B). k-mers not shared between both genomes (*Lk* for each genome) are shaded in grey. **(b) Influence of neighboring SNPs.** The sequence of each genome is split into k-mers of length *k*=5. Both genomes differ in 2 SNPs (colored in red in genome B) located within *k* base pairs. *p* means the distance between SNPs, and it is calculated as the difference between the nucleotide position of each SNP (in this case, *p*=3). k-mers not shared between both genomes are shaded and they correspond to 2*k*-*p*-1 k-mers for each genome. **(c) Influence of sequence insertion.** The sequence of each genome is split into k-mers of length *k*=5. Genome B is equal to genome A, except for a *L*=7 bp insertion (indicated in red letters). k-mers that are not shared by both genomes are shaded in grey, and they correspond to *k*-1 and *L*+*k*-1 k-mers, respectively. **(d) Influence of sequence deletion.** The sequence of each genome is split into k-mers of length *k*=5. Genome B is identical to genome A, except for a *L*=7 bp deletion. k-mers that are not shared between both genomes are shaded in grey, and they correspond to *L*+*k*-1 and *k*-1 k-mers, respectively. **(e) Influence of sequence replacement.** The sequence of each genome is split into k-mers of length *k*=5. Genome B has a substitution of *L*=7 bp respect to genome A, which is indicated in red letters. k-mers that are not shared between both genomes are shaded in grey, and they correspond to *L*+*k*-1 k-mers for each genome. **(f) Relationship between JI and %id.** Each curve represents the comparison of a reference genomes of 5 Mb with a second genome. *x*-axis represents both the percent identity (%*id*) between the genomes and the corresponding number of different base pairs between them. *y*-axis represents the Jaccard Index, using k-mers of 21 bp. The cases analyzed were: 1. The second genome differs from the reference only by SNPs allocated using a Poisson model, according to Eq. 6 (JI=0.21208 for id=0.95%); 2. The second genome differ only by SNPs, but the minimal distance between two SNPs is *k* base pairs, Eq. 5 (JI=0 for id=0.95%); 3. The second genome contains insertions, Eq. 8 (JI=0.95237 for id=0.95%); 4. The second genome differs by sequence deletions, Eq. 10 (JI=0.94999 for id=0.95%); and 5. The second genome differs by sequence substitutions, Eq. 11 (JI=0.90475 for id=0.95%). A discontinuous orange line indicates for case 2 the %id for which the minimal distance between two SNPs can no longer be *k* base pairs (id=0.95238%=1-(1/*k*)). A discontinuous magenta line indicates the JI value for which 5,880 random SNPs (id=0.99882%, case 1) are equivalent to a 250 kb sequence insertion (id=0.95%, case 3).

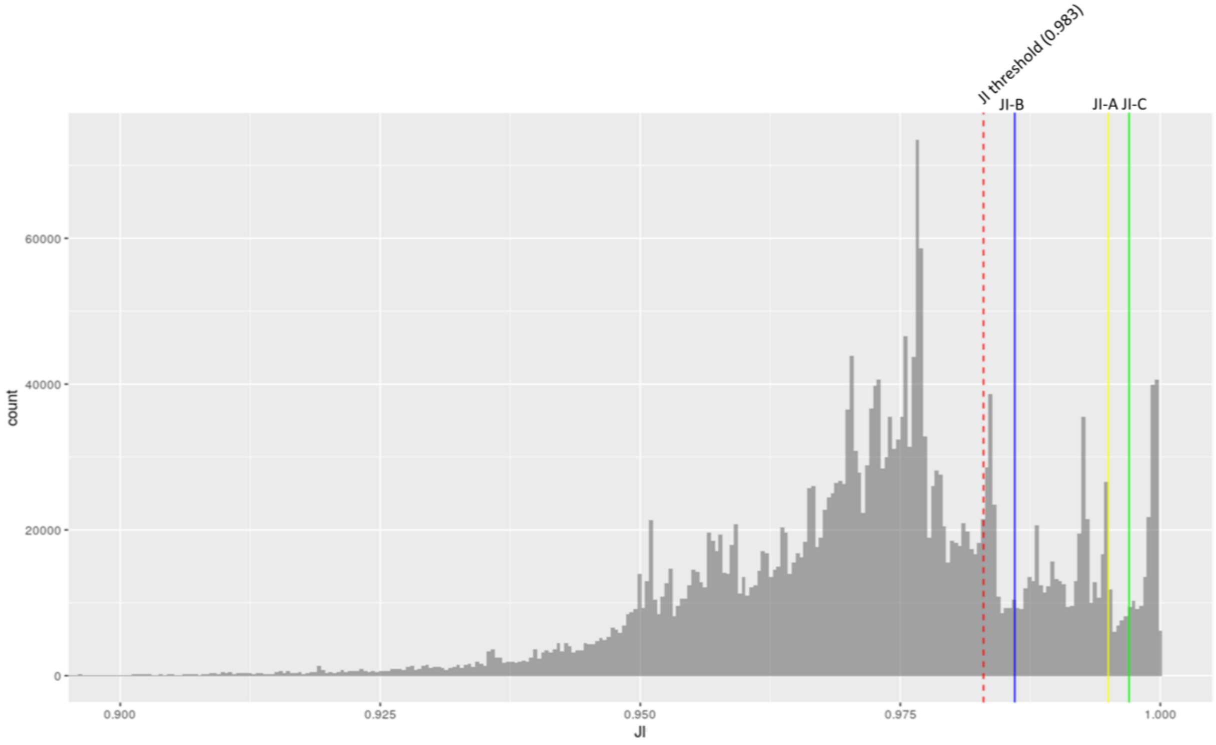


**Figure S2: JI distribution obtained from the pairwise comparison of *Salmonella* Typhi genomes.** Histogram of the JI values of pairwise Typhi genome comparisons. The JI threshold used in this study (JI=0.983) is indicated by a discontinuous red line. Vertical lines in different colors indicate the JI values selected to further analyze the within-groups differences for JI-A, JI-B and JI-C groups.

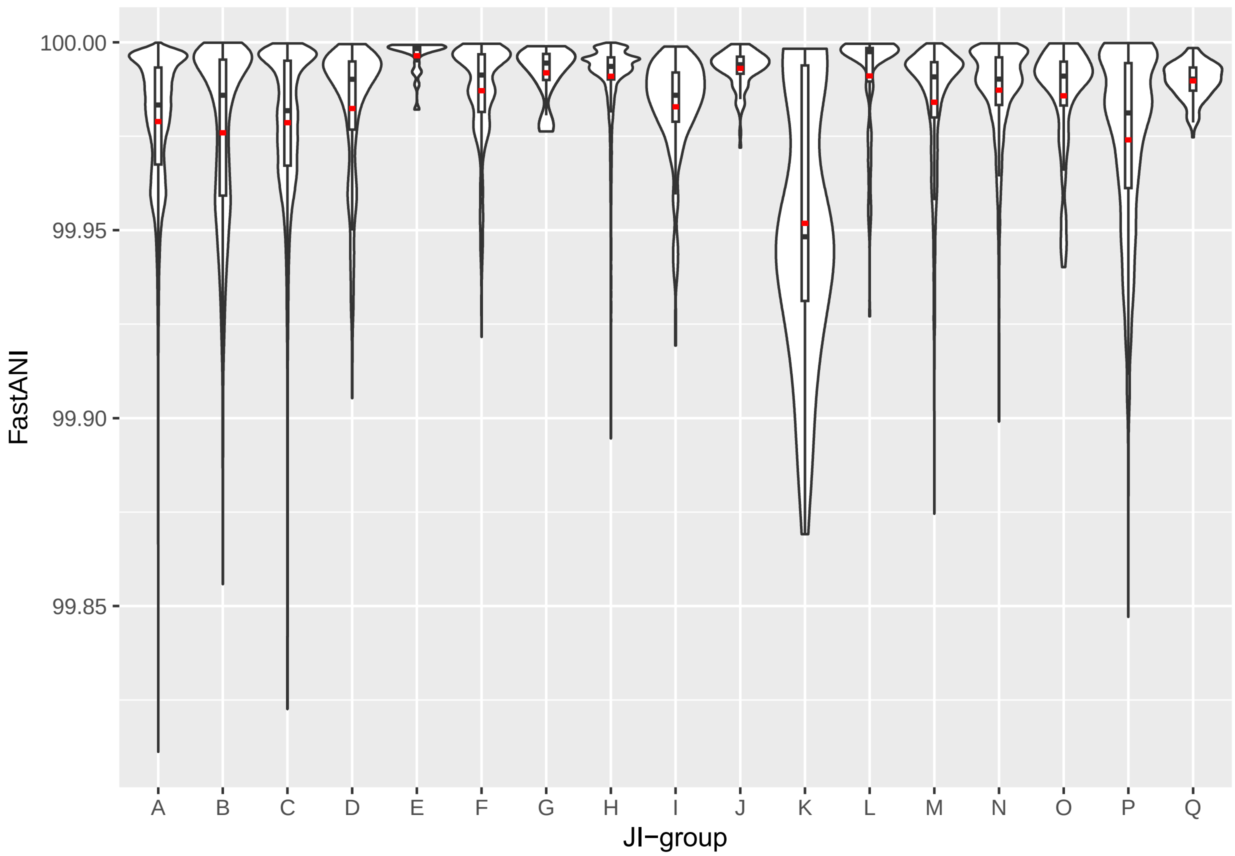


**Figure S3: Relatedness of genomes within each JI-group.** Violin plots showing the distribution of the FastANI intra-group pairwise genome comparisons for the 17 JI-groups, excluding those of each genome with itself. A boxplot is included into each violin, delimited by the first and third quartiles. Within each boxplot, horizontal lines represent the median (black) and the average (red) values.

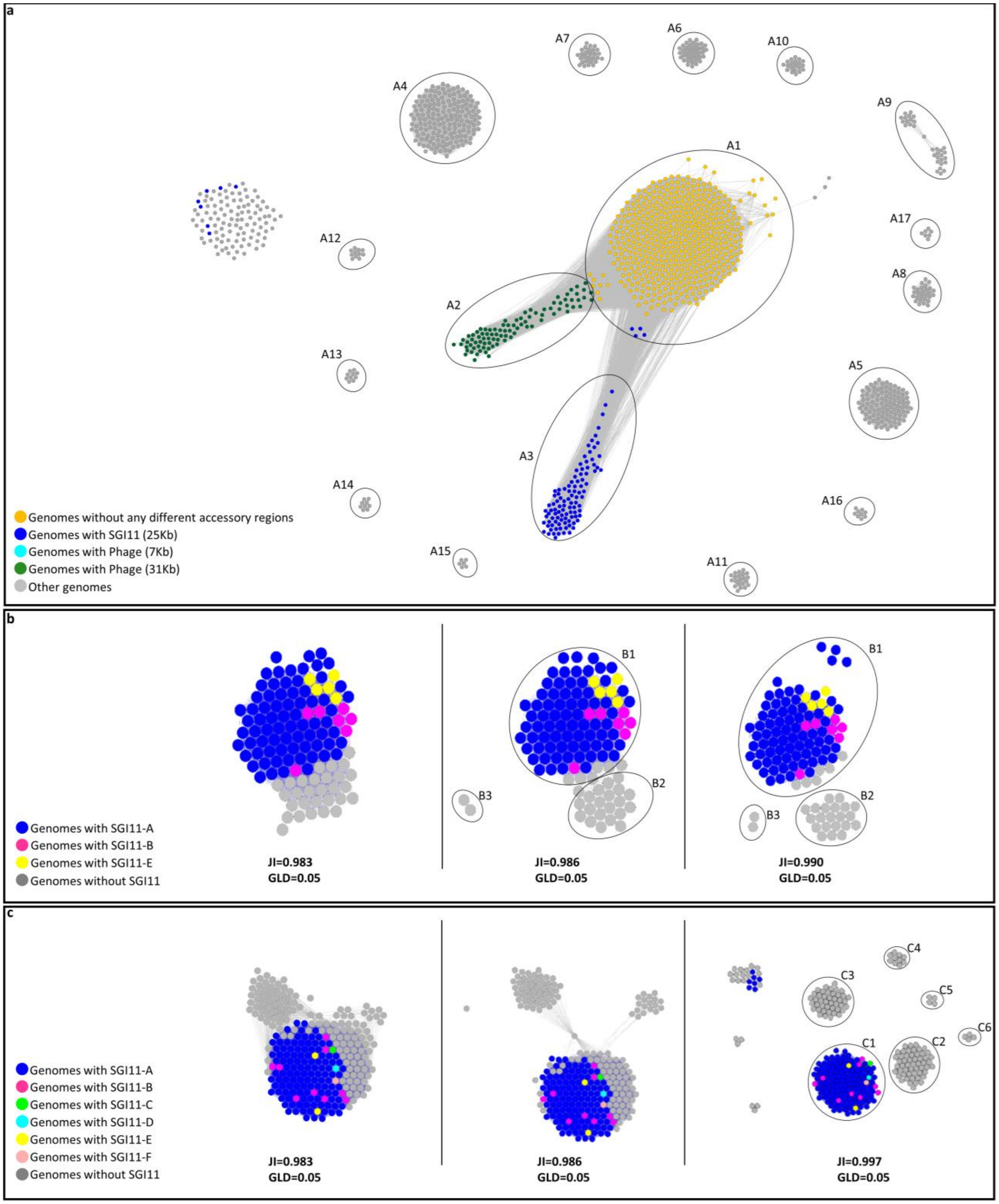


**Figure S4: Subclustering analysis of JI-groups A, B, and C. (a) Subclustering analysis of JI-Group A.** The network contains 1,319 genomes of JI-group A, using JI=0.995 as a threshold. Subgroups A1 to A17 defined by the Louvain method are surrounded by circles. Genomes of subgroups A2, A3, and A4 are colored by the presence/absence of SGI11 or other accessory regions. **(b) Subclustering analysis of JI-group B.** JI networks of 114 JI-B genomes at GLD=0.05 and increasing JI thresholds are shown. Subgroups determined by the Louvain method are indicated by circles. Nodes are colored according to the encoded SGI11 variant. **(c) Subclustering analysis of JI-group C.** JI networks of 265 JI-C genomes at

GLD=0.05 and increasing JI thresholds are shown. Subgroups determined by the Louvain method are indicated by circles when contain more than three genomes. Nodes are colored according to the encoded SGI11 variant. Membership of JI-A, JI-B and JI-C genomes to each subgroup can be found in column "JI-subgroup" of Supplementary Table S1.

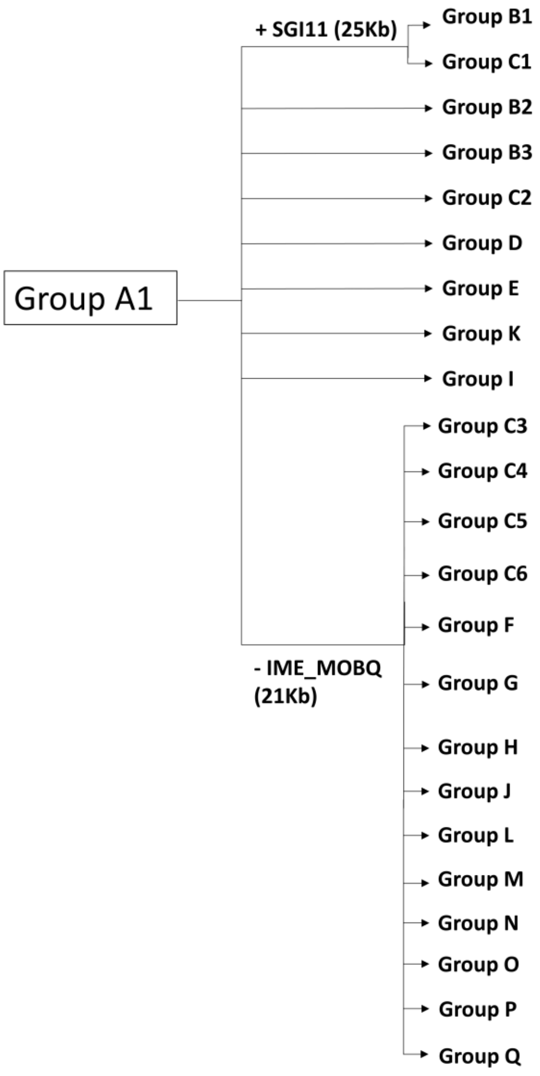


| | Lost | Gained |
|---|---|---|
| Group B1 | | PTU-E50 (83Kb) |
| Group C1 | | PTU-E18 (107Kb) |
| Group B2 | | PTU-E50 (93Kb) |
| Group B3 | | PTU-E50 (83Kb) |
| Group C2 | | PTU-E18 (107Kb) |
| Group D | | PTU-HI1A (216Kb) |
| Group E | | Region 1 (49Kb) |
| Group K | | PTU-Y (98Kb) |
| Group I | IME_MOBH (38 Kb), Region 2_MPFtype G_MPFtypeF (61Kb), Phage 1 (21Kb) | |
| Group C3 | Phage 2 (11Kb) | PTU-E18 (107Kb) |
| Group C4 | Region 3 (7Kb) | PTU-E18 (107Kb), Phage 3 (38Kb) |
| Group C5 | | PTU-E18 (107Kb), Phage 3 (38Kb), Phage 4 (33Kb), Region 4 (10Kb) |
| Group C6 | Region 3 (7Kb) | PTU-E18 (107Kb), Phage 3 (38Kb), Phage 4 (33Kb) |
| Group F | Region 5 (11Kb) | Region 6 (34Kb) |
| Group G | Region 2_MPFtype G_MPFtypeF (61Kb), Region 7 (62Kb), Phage 5 (16Kb), Region 8 (21Kb) | |
| Group H | | ICE_MOBH (55Kb) |
| Group J | | PTU-E50 (115Kb), Phage 6 (29Kb) |
| Group L | | Region 9 (67Kb) |
| Group M | | Phage 7 (50Kb) |
| Group N | Phage 1 (21Kb) | PTU-E80 (25Kb) |
| Group O | | Phage 8 (40Kb), Region 10 (40Kb) |
| Group P | | Phage 9 (44Kb) |
| Group Q | Region 5 (11Kb) | Region 11 (12Kb), Region 12 (11Kb) |

**Figure S5: Scheme of accessory genome dynamics by JI-groups.** Presence or absence of different accessory genome elements (plasmids, ICEs (integrative and conjugative elements), IMEs (integrative and mobilizable elements), phages, MPFs (mating pair formation systems, i.e, type IV secretion systems), and other genomic regions) in different JI-groups is indicated, taking JI-A1 subgroup as a reference.

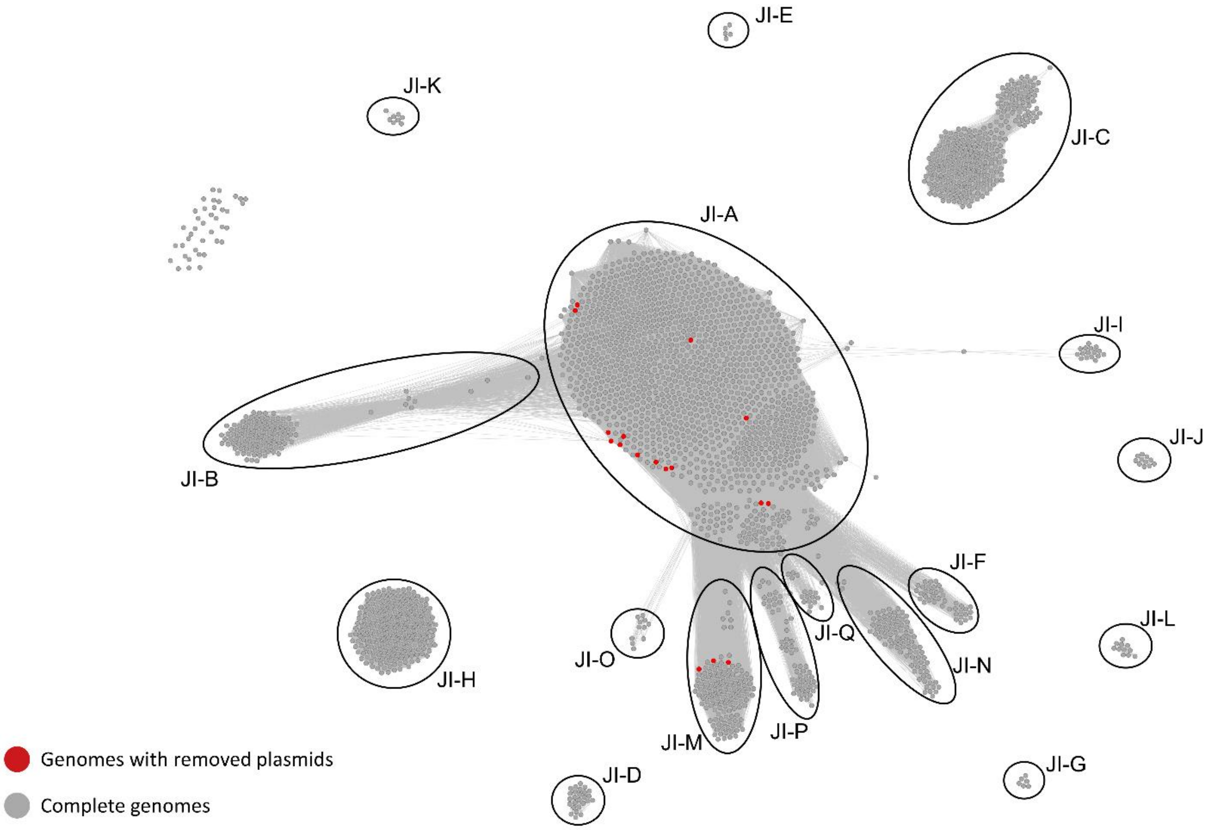


**Figure S6: Effect of plasmid removal on the JI network.** The plasmid sequences were removed from 13 RefSeq200 reference genomes present in JI-groups B, C, D, and K and 4 reference genomes of JI-groups C and J reconstructed by PLACNETw (column "Genomes with removed plasmids in Supp Fig S6" of Supplementary Table S1). The pairwise JI value was recalculated, and a new network generated. The "cured" genomes segregated from their original JI-groups (B, C, D, J, and K) and associated with the JI-A and JI-M genomes (in this latter case they come originally from JI-C). The network contains a total of 2,392 nodes, which are connected whenever JI = 0.983 and GLD = 0.05. Nodes colored in red (17) indicate reference genomes that contain plasmids larger than 25 kb in size, which were removed prior to JI calculation. Seventeen distinct clusters (named JI-A to JI-Q) identified by the Louvain method are indicated by circles.

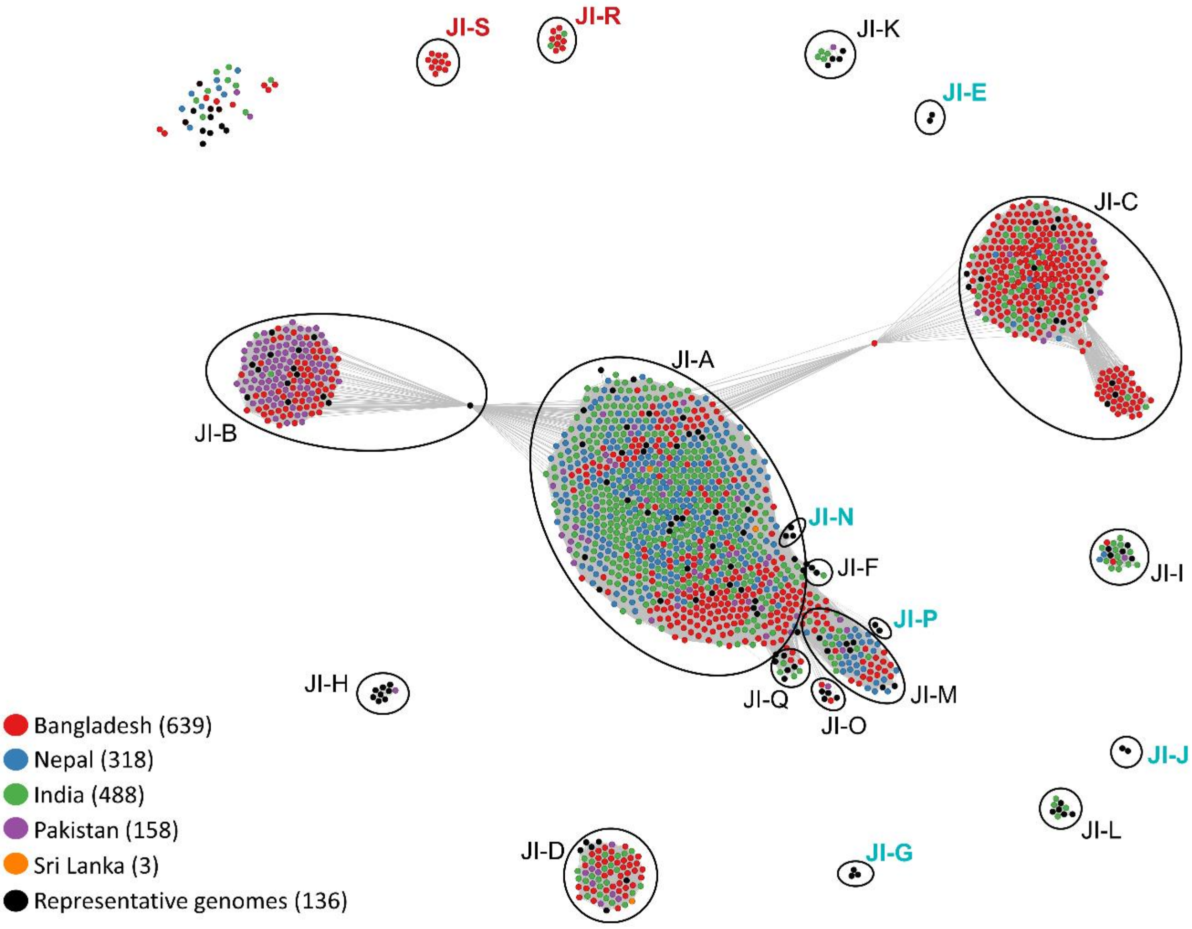


**Figure S7: Genomic diversity among *Salmonella* Typhi from the Indian subcontinent.** The network contains 1,606 Typhi genomes from the Indian subcontinent, and 136 reference genomes (from both US dataset and RefSeq200 as representatives of the different JI-groups) (Supplementary Table S2). Each genome is represented by a node, which is colored by its country of isolation as indicated in the legend. The number of isolates from each country is indicated between parentheses. Nodes are connected with thresholds JI=0.983 and GLD=0.05. Nineteen distinct clusters (JI-A to JI-S) detected by the Louvain method are indicated by circles. They are named in blue when only found in the representative genomes, in red when only present in the Indian subcontinent dataset, and in black when containing genomes from both datasets.

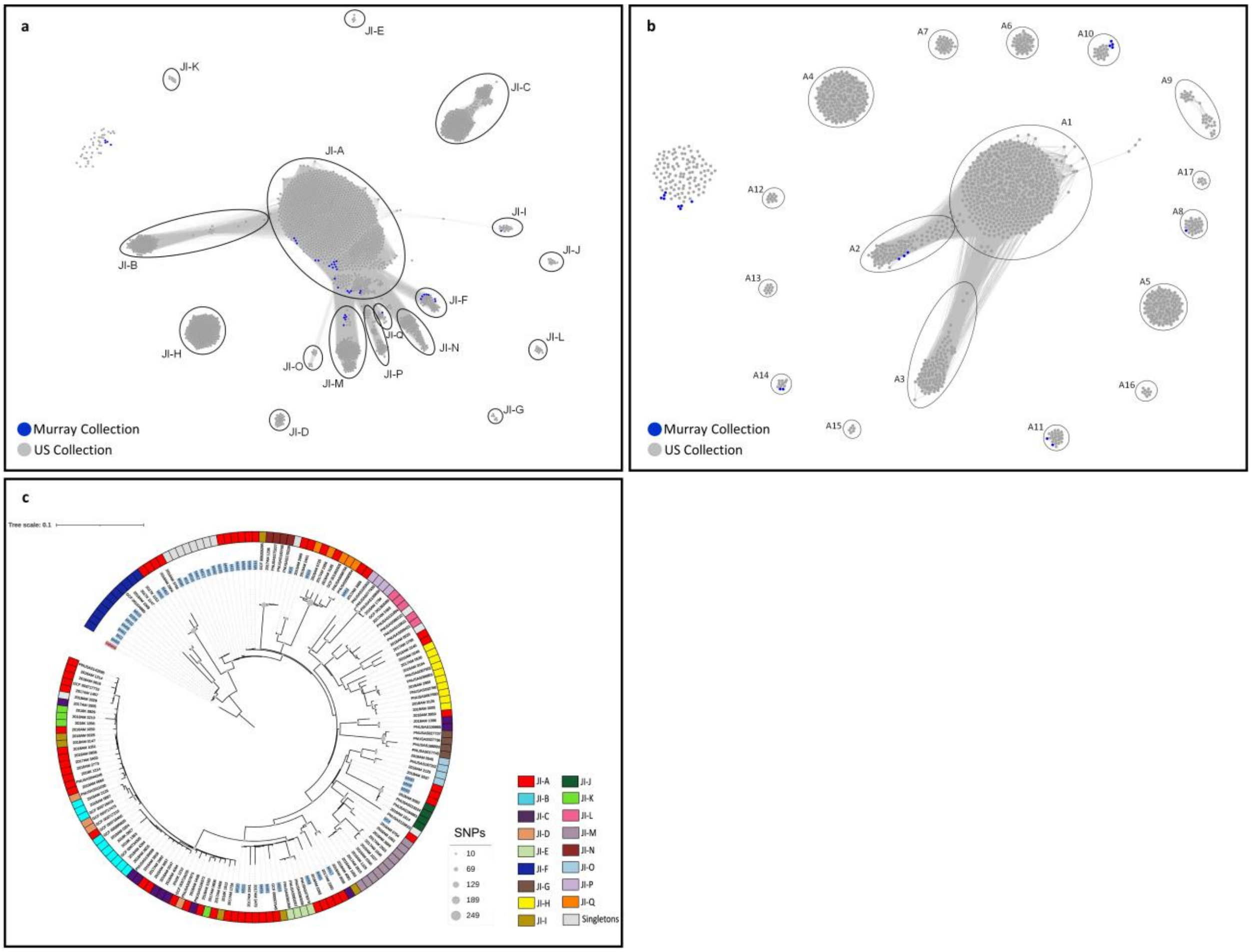


**Figure S8: Genomic diversity among *Salmonella* Typhi from the Murray collection. (a) Distribution of Typhi genomes of the Murray collection in the overall JI-network of Typhi**. The network contains all the Typhi genomes of the current study (n=2,392, colored in grey, Supplementary Table S1) and 38 Typhi genomes from the Murray collection, colored in blue (Supplementary Table S3). Nodes are connected whenever JI>=0.983 and GLD>=0.05. **(b) JI network of JI-A genomes.** The network contains 1,319 JI-A genomes (see column "JI-group" of Supplementary Table S1), and 20 JI-A genomes from the Murray collection (see column "JI-group" of Supplementary Table S3). This network was generated at JI=0.995. **(c) Phylogenetic distribution of Typhi genomes from the Murray collection.** Parsimony reconstruction based on the core SNPs using kSNP3.0 (*9*). The tree includes 130 representative genomes of the US dataset (see column "Genomes in Supp Figs S8c and S12" of Supplementary Table S1) and 38 Typhi genomes of the Murray collection (names labeled in blue). Genome NZ_CP015724.1 (serovar Indiana) was used as an outgroup and is labeled in red. The JI-group for each genome is indicated in the outer ring. Circles at the internal nodes indicate the number of SNPs that are shared exclusively by the descendands of each node. The tree was visualized with iTol v6 (*11*).

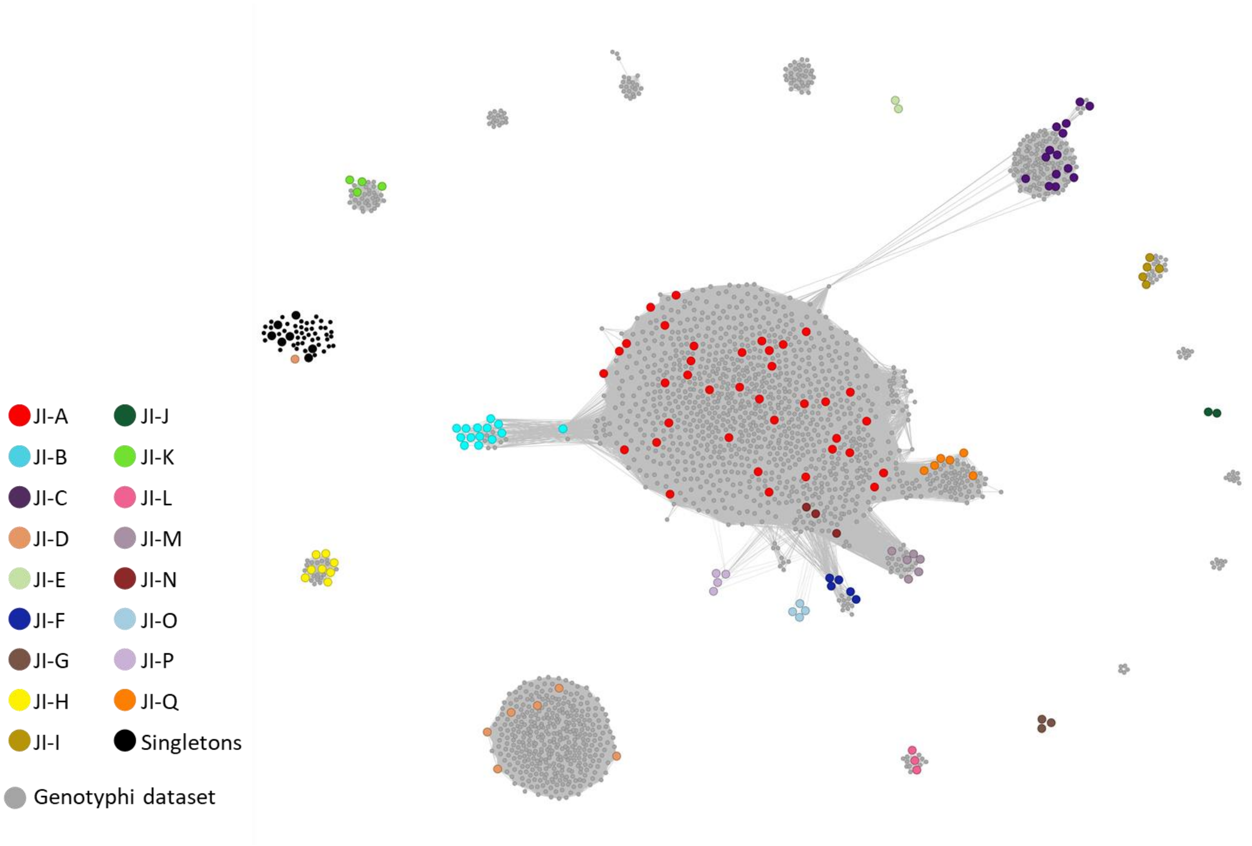


**Figure S9: Genomic diversity of globally representative Typhi genomes.** The network contains 1,804 Typhi genomes from the GenoTyphi dataset and 136 genomes (from both US dataset and RefSeq200 as representatives of the different JI-groups) (Supplementary Table S4). Nodes in the network represent genomes, with larger nodes corresponding to representative genomes, which are color-coded based on the JI-group they are assigned to. Nodes are connected with thresholds JI=0.983 and GLD=0.05.

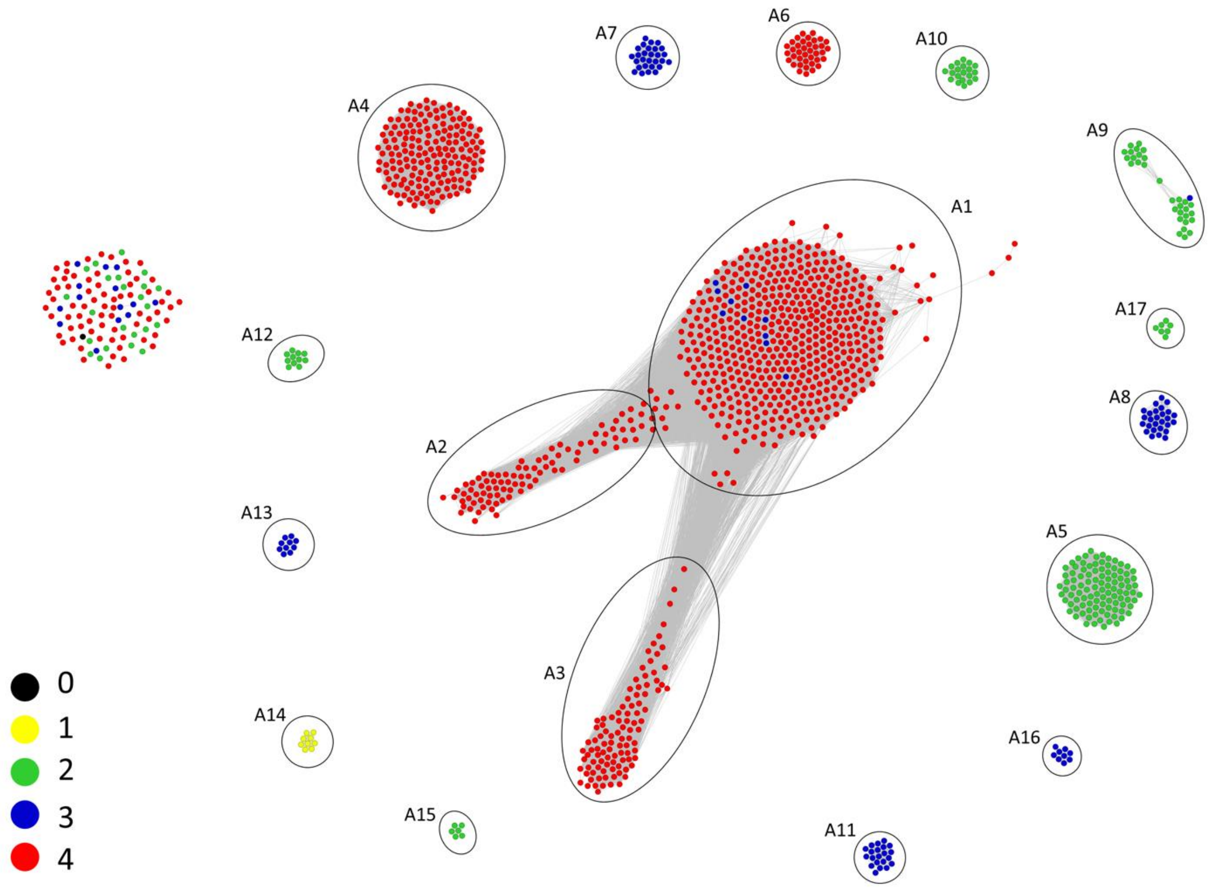


**Figure S10: Subclustering analysis of JI-Group A.** A set of 1,319 genomes of the JI-group A (Supplementary Table S1) were used to build the JI network, using JI=0.995 as a threshold. Subgroups A1 to A17 identified by the Louvain method are defined by circles. Nodes are colored by the GenoTyphi primary clade they belong to.

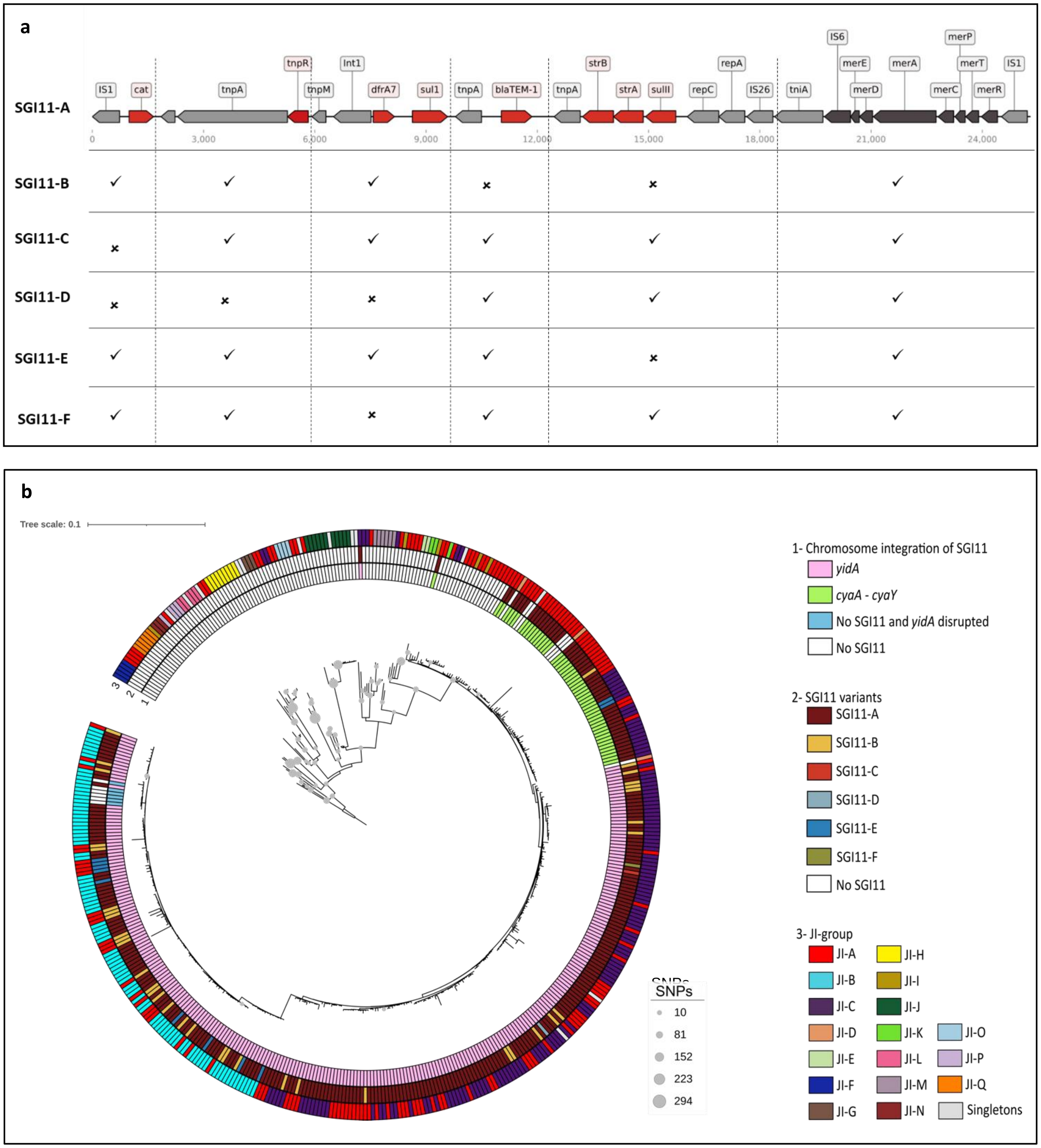


**Figure S11: SGI11 variants and their distribution in the Typhi genomes. (a) SGI11 variants found in the study genomes.** Genes encoded by the SGI11 variant A, present in *S*. Typhi BD1380 (GenBank Acc. No. KM023773), are represented by arrows and named in the upper part. They are colored in red (antimicrobial resistance genes), dark grey (*mer* operon), or light grey (other genes). In the lower part, a table records the presence or absence of different SGI11 segments in the different variants found in the study genomes. SGI11 variants A-E were previously described (*12*), while SGI11-F is a new variant described here. **(b) Phylogenetic distribution of SGI11.** Parsimony reconstruction based on the core

SNPs using kSNP3.0 (*9*) The tree includes 329 genomes that contain a chromosomal SGI11 insertion, 130 genomes that do not contain SGI11 (see column “Genomes in Supp FigS11” of Supplementary Table S1), and a non-Typhi genome (NZ_CP015724.1) that was used as an outgroup. Circles at the internal nodes indicate the SNPs shared exclusively by their descendants, according to the legend. Colored rings indicate the chromosomal SGI11 integration site (1) (see column “Chromosomal Insertions SG11 (ring 1 in Supp Fig S11)” in Supplementary Table S1), the SGI11 variant (2), and the JI-group of each genome (3). The tree was visualized with iTol v6 (*10*).

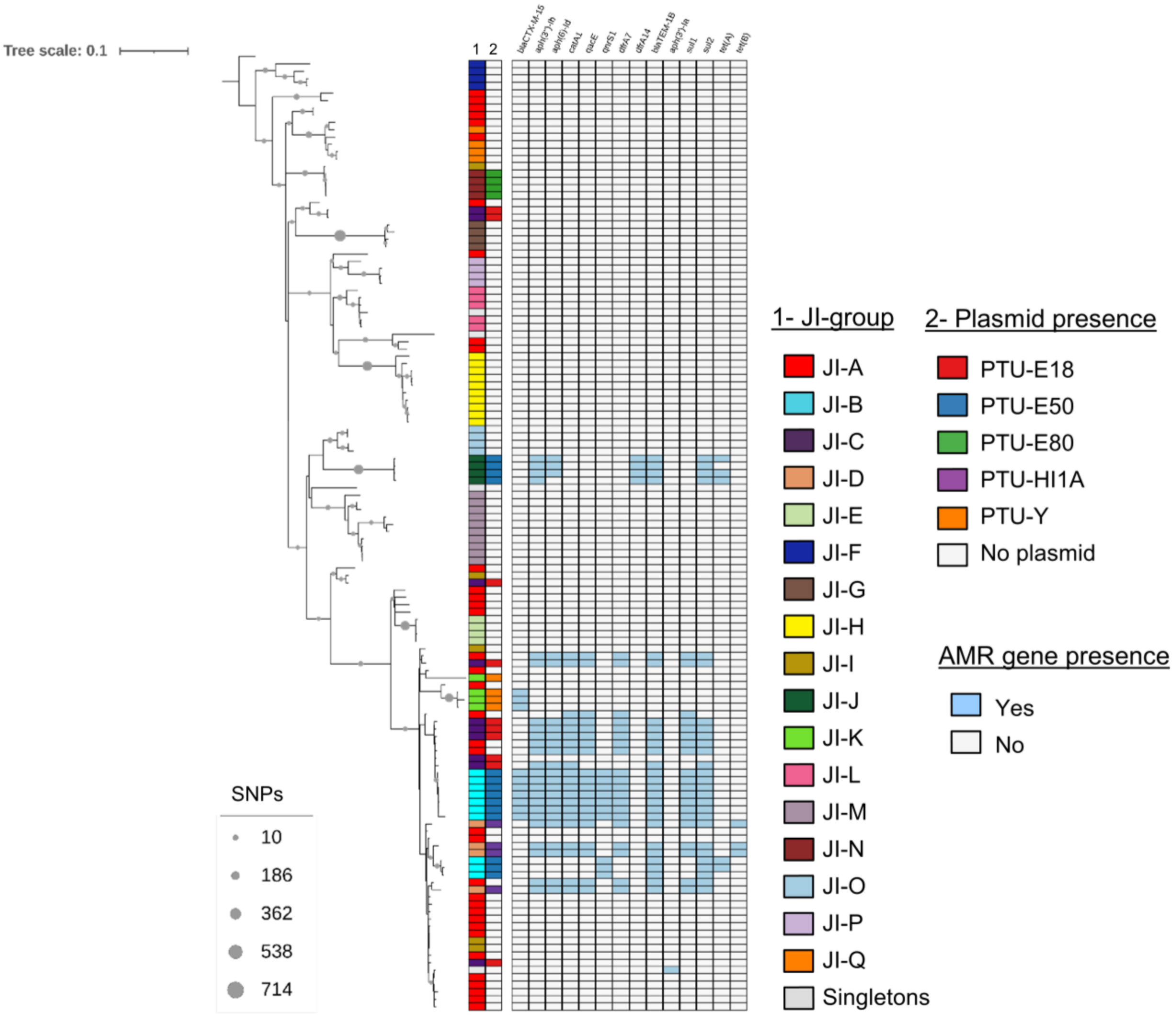


**Figure S12: Phylogenetic distribution of representative Typhi genomes.** Parsimony reconstruction based on the core SNPs using kSNP3.0 (*9*). The tree includes 130 Typhi genomes representatives from the 17 JI-groups of the US CDC dataset (see column "Genomes in Supp Figs S8c and S12" of Supplementary Table S1) and a genome from serovar Indiana (NZ_CP015724.1), which was used as outgroup. Circles at the internal nodes indicate the SNPs shared exclusively by their descendants, according to the legend. Colored rings indicate JI-group of each genome (1), the plasmid content (2), and AMR gene content. The tree was visualized with iTol v6 (*10*).

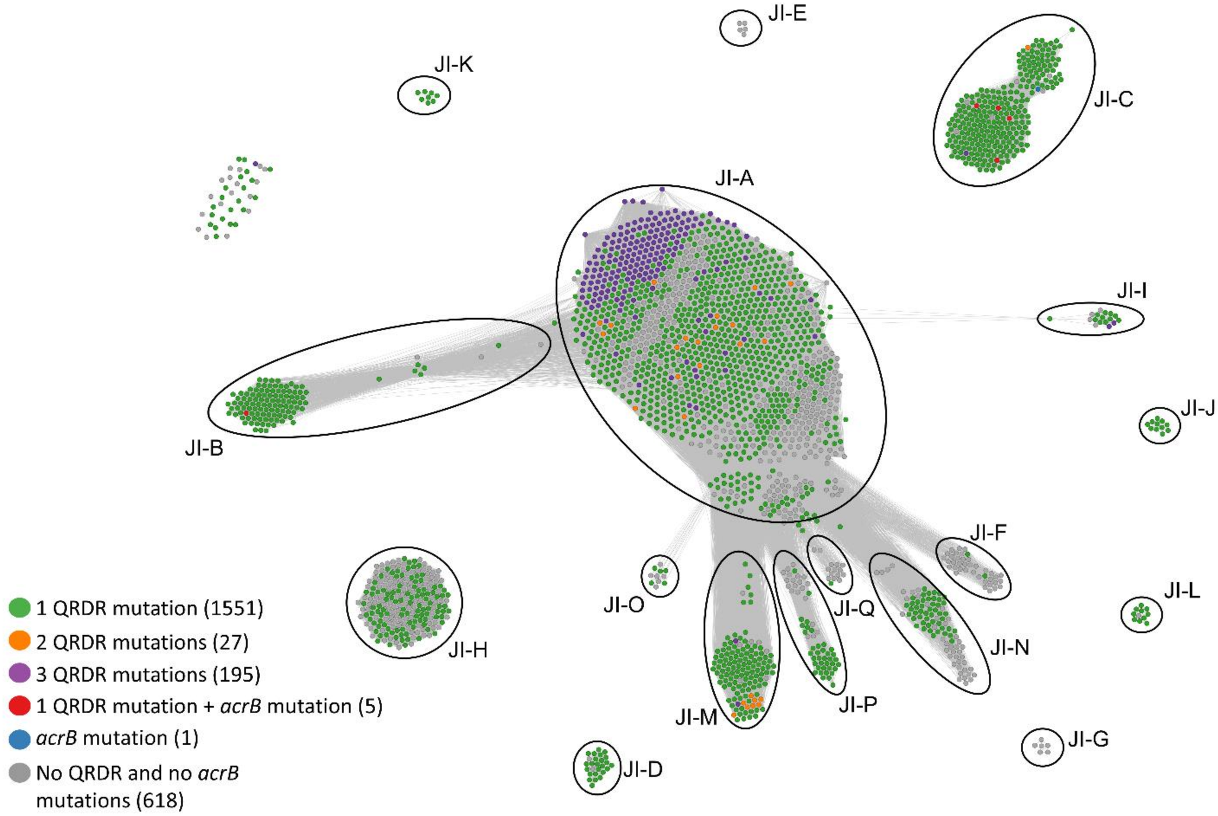


**Figure S13: Distribution of QRDR and *acrB* mutations in the JI network.** The network contains a total of 2,392 nodes that are connected whenever JI>=0.983 and GLD>=0.05. Seventeen distinct clusters (named JI-A to JI-Q) are indicated by circles. Nodes are colored according to the pattern of quinolone resistance determining region (QRDR; mutations in genes *gyrA*, *gyrB,* and *parC*), and the *acrB* gene mutations. The number of genomes with mutations are indicated at the left.

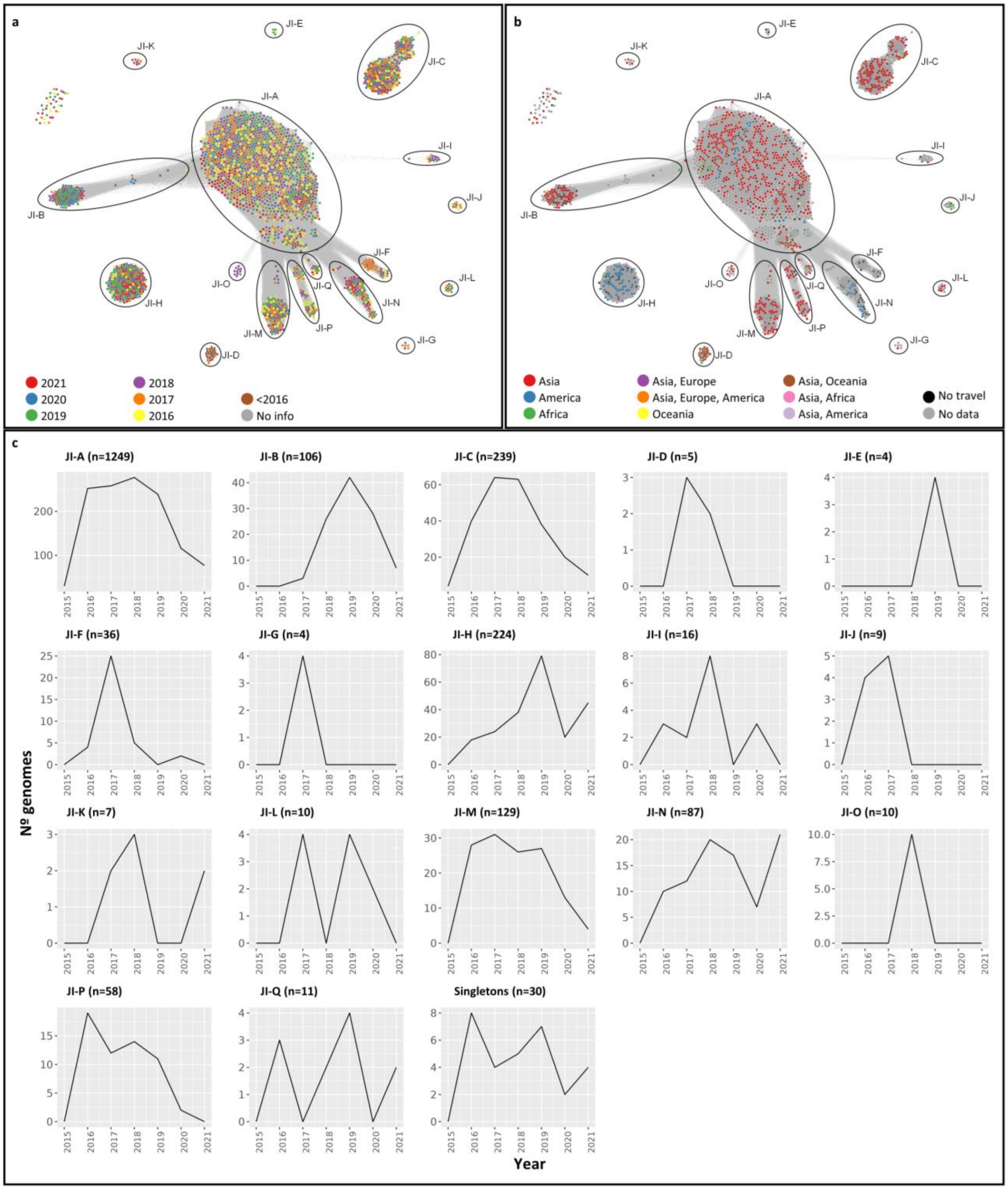


**Figure S14: Epidemiological data mapped onto the JI-network. (a) Temporal distribution of Typhi genomes in the JI-groups.** The network contains a total of 2,392 nodes (Supplementary Table S1) that are connected whenever JI>=0.983 and GLD>=0.05. Seventeen distinct clusters (JI-A to JI-Q) are indicated by circles. Nodes are colored according to the isolation year. **(b) Probable origin of the *S.* Typhi isolates.** Nodes are colored by United Nations Region of travel within 30 days of illness onset. **(c) Abundance of genomes of each JI-group over time.** Number of genomes of each JI-group during the period 2015-2021.

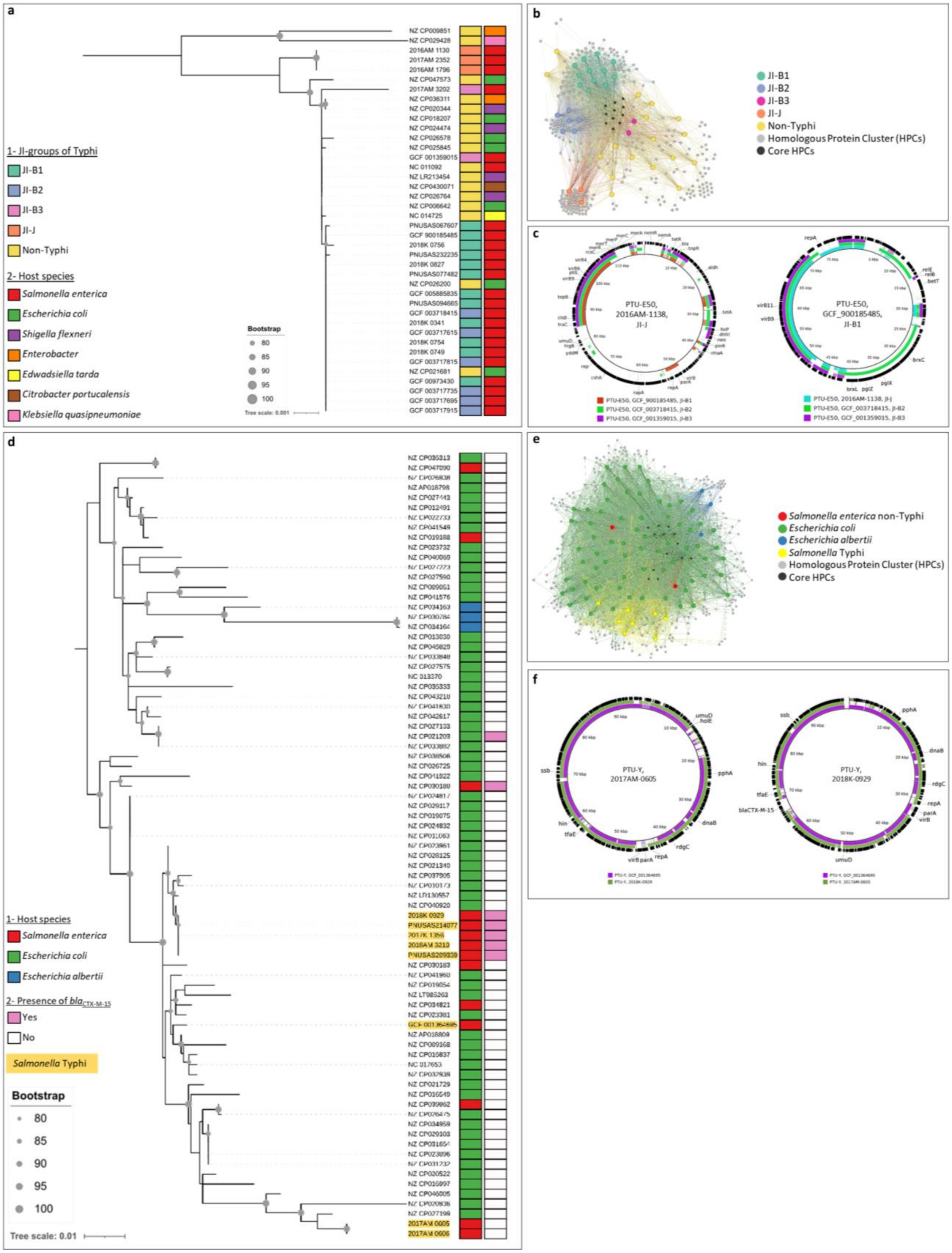


**Figure S15: Analysis of PTU-E50 and PTU-Y plasmids. (a) Core genome phylogenetic tree of PTU-E50 plasmids.** 23 plasmids present in *S*. Typhi (see PTU-E50 plasmids in column "PTU" of Supplementary

Table S1) and 17 plasmids from RefSeq200 non-Typhi *Enterobacterales* hosts were used to build a maximum likelihood (ML) tree based on the core genome by using IQ-TREE v2 (*3*). The tree was midpoint rooted and visualized with iTol v6 (*10*). UFBootstrap values > 80% are indicated by circles on the corresponding nodes. Close to the tips, colored strips indicate the JI-group (1) and the taxonomic species (2) of the plasmid host. **(b) Proteome network of PTU-E50 plasmids.** The proteins of the PTU-E50 plasmids were clustered at 80% identity and 80% coverage using AcCNET (*6*). The larger nodes correspond to plasmids and are colored according to the JI-group or subgroup of their bacterial host. The smaller nodes represent homologous protein clusters and are colored in black if contain members from all plasmids, or grey otherwise. Both kinds of nodes are connected if a plasmid contains a member in the corresponding protein cluster. **(c) Genome comparison of PTU-E50 plasmids.** In the left panel, a PTU-E50 plasmid of the JI-J group is used as reference and the colored rings represent regions of PTU-E50 plasmids from different JI-B subgroups shared with the reference. In the right panel, a PTU-E50 plasmid of the JI-B1 subgroup is used as reference and the colored rings represent regions of PTU-E50 plasmids from JI-J, JI-B2, and JI-B3 groups shared with the reference. In both cases, the outermost ring depicts the genes of the reference plasmid. The genomic comparisons were carried out with BRIG v0.95 (*8*). **(d) Core genome phylogenetic tree of PTU-Y plasmids.** 8 plasmids present in *S*. Typhi (see PTU-Y plasmids in column "PTU" of Supplementary Table S1) and 63 plasmids from RefSeq200 non-Typhi *Enterobacterales* hosts were used to build a ML tree as in **(a)**. Close to the tips, colored strips indicate the taxonomic species of the plasmid host (1) and the presence of $bla_{\text{CTX-M-15}}$ in the plasmid (2). The 8 plasmids from *S*. Typhi (JI-group K) are shadowed in yellow. **(e) Proteome network of PTU-Y plasmids.** The network was constructed as indicated in **(b)**. **(f) Genome comparison of PTU-Y plasmids.** Plasmids containing or not $bla_{\text{CTX-M-15}}$ are compared using BRIG (*8*). At the right, a PTU containing $bla_{\text{CTX-M-15}}$ is used as a reference and the colored rings represent regions of PTU-Y plasmids that do not encode this gene. At the left, a non-encoding $bla_{\text{CTX-M-15}}$ plasmid is used as a reference and the colored rings represent regions of PTU-Y plasmids that do not encode (inner ring) or encode this gene (outer ring). In both cases, the outermost ring depicts the genes of the reference plasmid.

**Dataset S1 (separate file):** US CDC and RefSeq200 dataset

**Dataset S2 (separate file):** Indian subcontinent dataset

**Dataset S3 (separate file):** Murray collection dataset

**Dataset S4 (separate file):** GenoTyphi collection dataset
